# EMMIs: Engineered Myometrial Microtissues for Direct Quantification of Oxytocin-Induced Contractility

**DOI:** 10.64898/2026.04.27.721112

**Authors:** Karla I Ortega Sandoval, Ritu M Dave, Cailin R Gonyea, Kaci Mitchum, Alana Aristimuno Millan, Shriya Suryakumar, Antonina I. Frolova, Shreya A Raghavan

## Abstract

Forceful and coordinated contractions of the uterine myometrium are essential for successful labor, delivery, and postpartum uterine involution. Failure of the uterus to generate or sustain contractile force (uterine atony) after delivery results in postpartum hemorrhage, a leading cause of maternal mortality globally. Paradoxically, uterine atony is exacerbated by prolonged oxytocin exposure used to induce or augment labor through a process of contractile desensitization. Despite its prevalent use in obstetrics, the direct impact of oxytocin desensitization on myometrial contractile force generation remains poorly defined. Current model systems are inadequate to address this gap: *ex vivo* myometrial tissue strips are limited by tissue availability, donor variability, and lack of genetic tractability, while existing *in vitro* models provide only indirect readouts of contractility without direct force quantification. Here, we introduce engineered myometrial microtissues (EMMIs), a platform enabling the direct, isometric measurement of contractile force in response to physiological agonists like oxytocin. By embedding and molding immortalized human myometrial smooth muscle cells within a collagen hydrogel, we induced significant structural and molecular maturation over six days. Upon maturation, EMMIs were characterized by circumferential cellular alignment, sustained expression of smoothelin, upregulation of connexin-43, and a transcriptomic shift toward a contractile phenotype. Mature EMMIs generated calcium-sensitive, dose-dependent contractions to oxytocin and potassium chloride. Genetic deletion of the oxytocin receptor abolished oxytocin-induced contractility, establishing receptor specificity. Finally, we utilized EMMIs to recapitulate clinical oxytocin desensitization, providing a direct link between prolonged oxytocin exposure and diminished contractile output. Together, these findings establish engineered myometrial microtissues (EMMIs) as a genetically manipulable, and reproducible system for investigating myometrial contractile physiology to improve obstetric outcomes.

**Teaser:** Engineered 3D uterine tissues quantify how labor-inducing drugs weaken contractions and drive maternal hemorrhage

## 1. Introduction

Myometrial contractility, *i.e.* the forceful and coordinated contractions of uterine muscle, is central to successful obstetric outcomes. Aberrant myometrial contractility underlies major obstetric complications, including labor dystocia, and life-threatening postpartum hemorrhage (PPH)(*1–3*). Failure of the uterus to contract effectively after delivery, a condition known as uterine atony, is the leading cause of PPH. The risk for uterine atony and hemorrhage increases with prolonged or repeated oxytocin exposure, which is widely used to induce, sustain, and augment labor(*4–6*).

Prolonged or repeated oxytocin exposure induces desensitization of the oxytocin receptor (OXTR). This process is documented both *ex vivo* and *in vitro* through reduced receptor binding and impaired downstream signaling, leading to attenuated myometrial responses to subsequent oxytocin stimulation(*7, 8*). Clinically, prolonged oxytocin exposure-induced desensitization compromises the uterus’s ability to sustain coordinated contractions after delivery, leading to uterine atony and PPH(*4–6*). Despite this well-established mechanistic sequence, existing model systems lack tissue-level complexity, leaving a critical gap: the direct impact of prolonged oxytocin exposure on functional contractile force generation remains poorly defined. This gap directly contributes to our inability to predict and prevent uterine atony, and therefore postpartum hemorrhage.

In this study, we introduce human, engineered myometrial microtissues (EMMIs) that tackles this gap by advancing beyond traditional *in vitro* and *ex vivo* methods of assessing myometrial contractility. Human myometrial tissue strips obtained at cesarean delivery represent the current gold standard for *ex vivo* functional assessment of uterine contractility and have been instrumental in characterizing responses to oxytocin, prostaglandins, and other uterotonic agents(*9, 10*). Organ bath studies using pregnant human myometrial strips have demonstrated that continuous or repeated oxytocin pretreatment attenuates subsequent oxytocin-induced contractions, consistent with receptor desensitization and signaling exhaustion(*11, 12*). However, strip-based *ex vivo* approaches are limited by restricted tissue availability, short-term viability in culture, high donor variability, and heterogeneous cellular composition, including variable proportions of smooth muscle cells, fibroblasts, immune cells, and vasculature. These factors hinder mechanistic dissection, complicate dose–response comparisons and impede experimental reproducibility across studies(*13–15*).

Isolated primary human myometrial smooth muscle cells or telomerase-immortalized human myometrial (hTERT-HM) cell lines have been developed to study contractility *in vitro*(*16*). These cells retain key contractile features, including expression of smoothelin and other contractile apparatus proteins critical for force generation, connexin-43 gap junctions that enable electrical and metabolic coupling, and functional oxytocin receptors that link hormonal stimulation to intracellular calcium elevation and downstream contraction(*16–19*). These properties make myometrial cell cultures attractive tools for dissecting signaling pathways. However, most existing *in vitro* assays provide only indirect readouts of contractility and do not directly quantify force output. For example, collagen gel contraction assays quantify matrix compaction as a composite measure of both cell-generated force and extracellular matrix (ECM) remodeling, but they cannot readily decouple these two processes or translate compaction into calibrated force values(*20–22*). Calcium imaging tracks intracellular signaling but reports only upstream calcium dynamics rather than mechanical output, and changes in calcium transients do not always correspond linearly to tissue-level force(*23*). Although recent three-dimensional systems such as bioprinted rings and microfluidic co-culture platforms integrate tissue architecture, none directly quantify isometric contractile force, which remains the physiological gold standard established by tissue strip studies(*14, 24, 25*). This gap is significant: without direct force quantification, the functional consequences of oxytocin receptor desensitization, and other pathophysiological factors, on contractile output cannot be resolved in a genetically tractable, reproducible system.

Here, we introduce engineered myometrial microtissues (EMMIs): a functional three-dimensional *in vitro* platform that bridges the gap between cell cultures and tissue strips by enabling the direct quantification of isometric, contractile force in a scalable human model. By integrating force measurements with genetic tractability, EMMIs are a robust platform to resolve functional consequences of oxytocin receptor desensitization, a primary driver of postpartum hemorrhage. Our findings demonstrate that EMMIs not only recapitulate complex myometrial physiology but are a powerful discovery platform for mechanistic and therapeutic targets in labor-related pathologies.

## 2. RESULTS

### 2.1 Bioengineering and Maturation of Engineered Myometrial Microtissues (EMMIs)

Immortalized human myometrial smooth muscle cells (hTERT-HM) were encapsulated within collagen I hydrogels and placed around a central post defining the hollow lumen. A schematic of the process is depicted in **Figure 1A**. Representative brightfield photomicrographs of engineered myometrial microtissues captured daily from day 0 to day 6 are shown in **Figure 1B**, demonstrating progressive compaction of the collagen hydrogel around the central post over the culture period. Quantification of gel area over time revealed that the majority of compaction occurred between day 0 and day 1, with a 90.78% reduction in gel area, followed by a gradual plateau through day 6 (**Figure 1C**). Wet weight measurements at days 1, 3, and 6 demonstrated a significant reduction of 68.80% from day 1 to day 3, stabilizing thereafter, consistent with ongoing matrix remodeling and fluid redistribution during early compaction (**Figure 1D**; n≥6; ****p<0.0001; one-way ANOVA).

**Figure 1.**
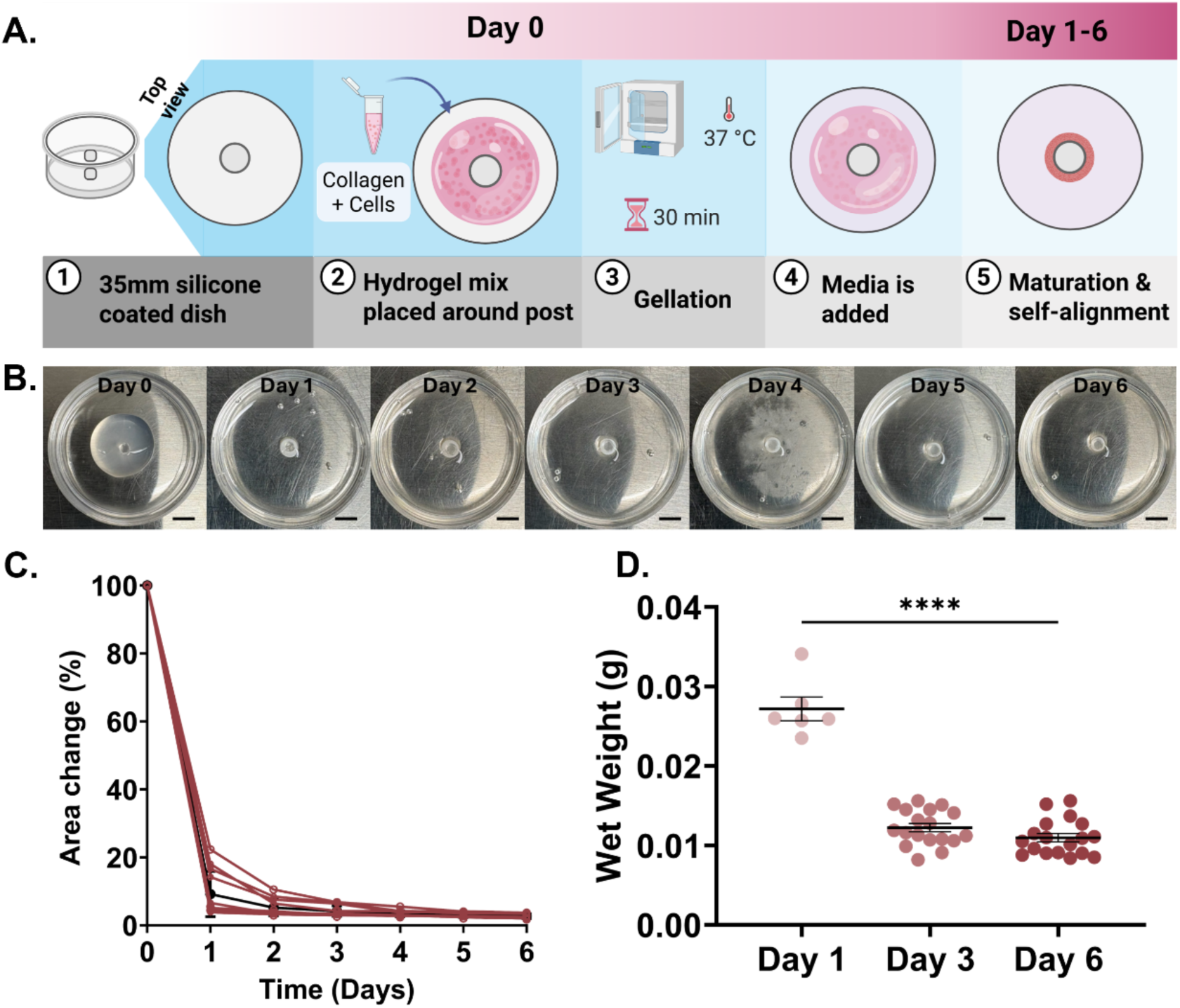
Fabrication of Engineered Myometrial Microtissues (EMMIs) (A) Schematic of the microtissue fabrication workflow. A hydrogel mixture of collagen type I and hTERT-HM cells was cast around a central silicone post in a 35 mm silicone-coated dish to define a hollow luminal space. Following gelation at 37°C for 30 minutes, culture medium was added, and constructs were maintained for up to 6 days to permit self-assembly into a ring-like geometry. (B) Representative photomicrographs of engineered myometrial microtissues captured daily from day 0 to day 6, showing progressive compaction and self-alignment around the central post. Scale bar = 5 mm. (C) Quantification of microtissue area change (%) over the 6-day maturation period, demonstrating progressive compaction, over 10 replicates of EMMIs. (D) Wet weight of microtissues at days 1, 3, and 6 showing significant reduction and stabilization of wet weight across 6 days (****p<0.0001, n ≥ 6, one-way ANOVA).

### 2.2 Structural and Molecular Maturation of EMMI

To characterize the structural and molecular maturation of engineered myometrial microtissues over time, whole-mount immunofluorescence imaging and quantitative western blot analysis were performed at days 1, 3, and 6 of culture. Together, these complementary approaches revealed progressive, time-dependent changes in cellular architecture, organization, and phenotypic maturation (**Figure 2A**).

**Figure 2.**
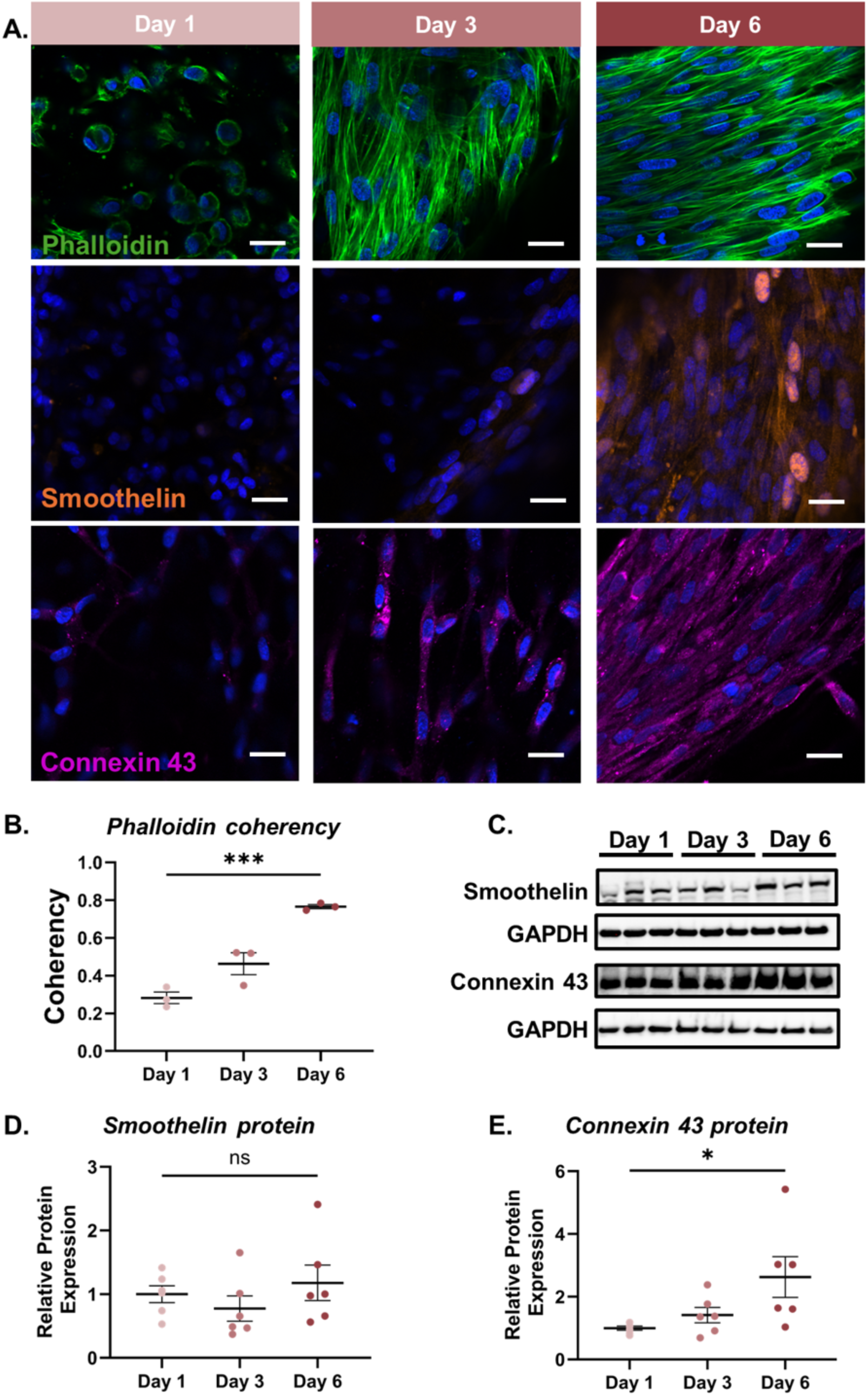
Characterization of EMMI Structural Organization and Maturity. (A) Representative whole-mount immunofluorescence images of EMMIs at days 1, 3, and 6, stained for F-actin (phalloidin, green), smoothelin (orange), and connexin-43 (magenta). Nuclei are counterstained with DAPI (blue). Scale bar = 20 µm. (B) Quantification of cytoskeletal coherency across maturation timepoints, indicating progressively increasing cell alignment (***p<0.001; n=3; one-way ANOVA); (C) Representative western blots for smoothelin (110 kDa) and connexin-43 (43 kDa), with GAPDH (36 kDa) as a loading control, at days 1, 3, and 6. (D) Densitometric quantification of smoothelin expression normalized to GAPDH across culture time, demonstrating slight non-significant increases (ns, not significant; n=6, one-way ANOVA). (E) Densitometric quantification of connexin-43 expression normalized to GAPDH across culture time, demonstrating significant increases in connectivity with maturation (*p<0.05, n=6, one-way ANOVA).

Assessment of cytoskeletal architecture via phalloidin staining demonstrated marked changes in cell morphology and actin organization over the culture period. At day 1, cells exhibited predominantly rounded morphologies with minimal elongation or directional alignment, and actin filaments appeared largely disorganized. By days 3 and 6, cells adopted elongated, spindle-shaped morphologies and assembled into circumferentially aligned networks along the ring axis, indicative of self-organization into smooth muscle-like fascicles (**Figure 2A**). Cytoskeletal alignment was quantified by measuring actin fiber coherency using OrientationJ (an ImageJ plugin), where a coherency value of 0 indicates randomly oriented fibers with no directional preference, and a value of 1 indicates complete uniaxial alignment. Coherency values in EMMIs increase as cells transition from a disorganized arrangement toward a highly aligned organization. Coherency increased significantly over time, from 0.28 ± 0.03 at day 1 to 0.77 ± 0.01 at day 6, representing a 2.71-fold increase relative to day 1 (***p<0.001, n=3; one-way ANOVA; **Figure 2B**).

Concurrent with these morphological changes, immunofluorescence imaging revealed progressive changes in the spatial distribution and organization of smoothelin over the culture period, with signal appearing increasingly abundant and localized along elongated cells by day 6(*26, 27*).

In parallel, the expression of connexin-43 (Cx43), a gap junction protein critical for intercellular electrical and metabolic coupling, was evaluated across culture timepoints. At day 1, Cx43 immunofluorescence signal was sparse and discontinuous, indicative of limited intercellular connectivity. With advancing maturation, Cx43 expression became markedly more abundant and exhibited a continuous distribution along cell borders, consistent with the progressive establishment of functional gap junctions within the three-dimensional tissue architecture.

Protein expression was quantified via western blot analyses, with representative blots shown in **Figure 2C**. Smoothelin protein levels remained stable over the culture period (1.18-fold; ns, n=3; one-way ANOVA; **Figure 2D**) and Cx43 protein levels increased over the same period (2.63-fold; *p<0.05, n=3; one-way ANOVA; **Figure 2E**).

### 2.3 Unbiased examination of EMMI maturation via transcriptomic characterization

To further characterize the maturation of EMMIs over time, RNA was extracted from day 1, 3, and 6 microtissues, and bulk RNA sequencing was performed. Principal component analysis demonstrated clear separation of samples by culture day, with clustering at each time point (**Figure 3A**). PC1 captured 84% of the total variance and primarily distinguished day 6 from day 1 and 3, while PC2 accounted for 10% and separated day 1 from day 3. This indicated that the transition from day 1 to day 3 and the transition from day 3 to day 6 represent two sequential, but not equal, phases of transcriptional change.

**Figure 3.**
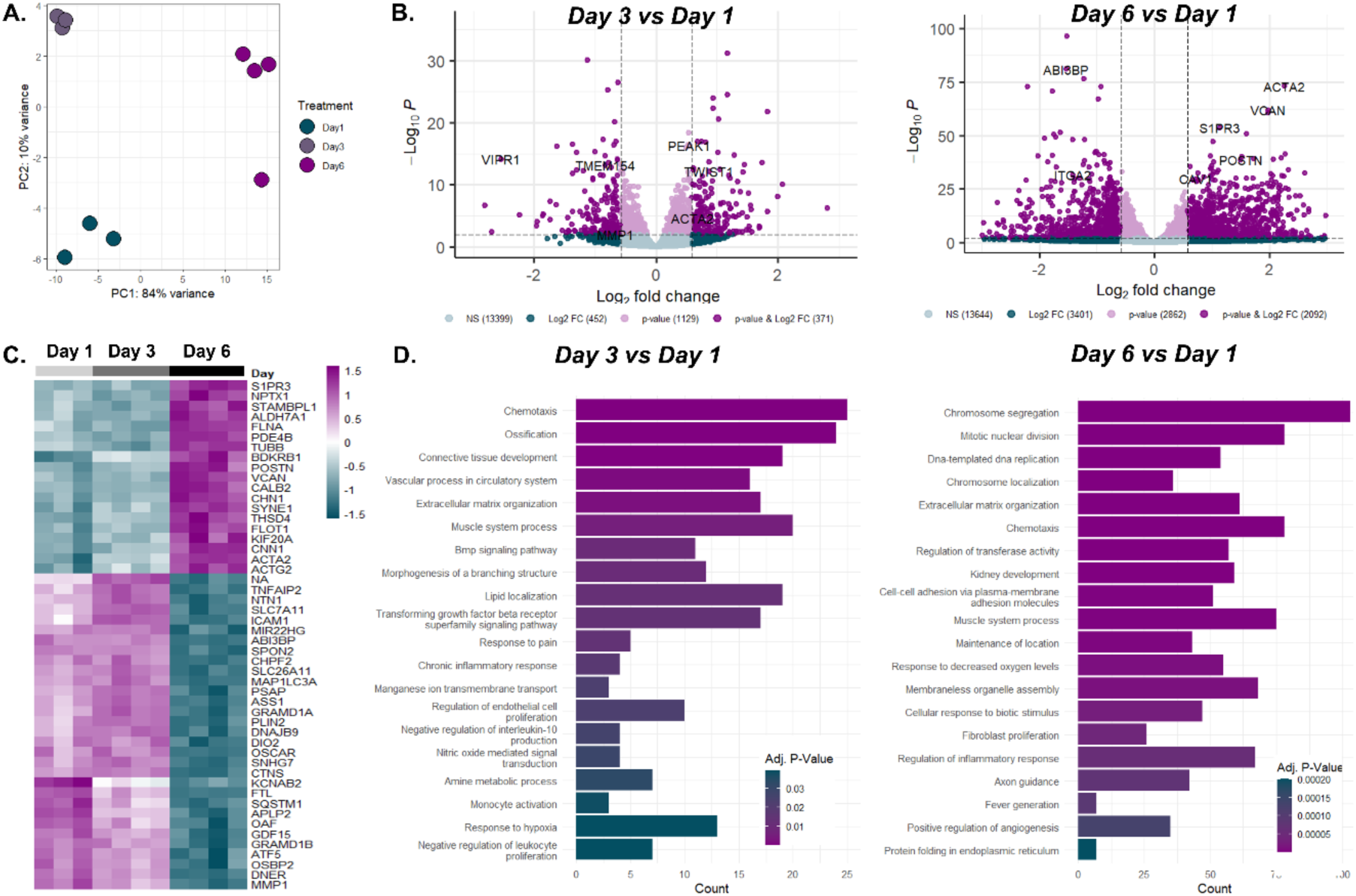
Transcriptomic Maturation of Engineered Myometrial Microtissues. (A) PCA plot of variance-stabilized counts for day 1, 3, and 6 microtissues (B) Volcano plots showing differentially expressed genes (DEGs) when comparing day 6 or day 3 to day 1 microtissues. Cutoff values of p adjusted for multiple comparisons (padj) < 0.01 and log_2_fold expression change > 0.58 were used to define significance. (C) Hierarchical clustering of the top 50 DEGs across day 1, 3, and 6 microtissues. (D) Top twenty enriched biological process pathways identified using clusterProfiler.

Differential expression analysis (padj < 0.01; log₂ fold change > 0.58) identified 371 DEGs between day 3 and day 1 microtissues, and 2,092 DEGs between day 6 and day 1, reflecting progressive and extensive transcriptomic remodeling over the culture period (**Figure 3B**). By day 3, the smooth muscle contractile marker α-smooth muscle actin (*ACTA2*) and the mesenchymal transcription factor *TWIST1* were among the upregulated transcripts, while the relaxant GPCR *VIPR1* was significantly downregulated. Together, these early shifts signal initiation of the contractile program alongside suppression of relaxant signaling and matrix remodeling. By day 6, *ACTA2* induction had intensified and extended into a broader smooth muscle and matrix signature, with coordinated upregulation of the pro-contractile GPCR *S1PR3*, the membrane scaffold *CAV1*, and the structural matrix components *POSTN* and *VCAN*(*28*).

Hierarchical clustering of the top fifty DEGs grouped day 1 and day 3 samples, separate from day 6 (**Figure 3C**). The day-6 cluster was dominated by contractile and ECM transcripts including the smooth muscle actins ACTA2 and ACTG2, the actin-binding regulator calponin (CNN1), the actin crosslinker filamin A (FLNA), the contraction-coupled GPCRs *BDKRB1* and *S1PR3*, and the structural matrix gene *POSTN*, confirming that the majority of transcriptomic maturation, particularly of the contractile program, occurs between days 3 and 6.

Gene Ontology enrichment analysis of the DEGs from each comparison identified the biological processes most altered during maturation (**Figure 3D**). At day 3, enriched terms included chemotaxis, extracellular matrix organization, muscle system process, response to hypoxia, and chronic inflammatory response, reflecting an early adaptive response as freshly compacted tissues recover from seeding while initiating contractile and structural programs. By day 6, enriched terms shifted toward chromosome segregation, DNA replication, muscle contraction, and cell-cell adhesion, alongside regulation of inflammatory response, indicating that microtissues continued to expand, contractile program matured, and the acute post-fabrication stress response moved towards resolution.

### 2.4 Time-dependent acquisition of contractile maturity and function in Engineered Myometrial Microtissues

Having established progressive structural and molecular maturation, functional contractility was subsequently assessed via isometric force measurements at days 3 and 6 in response to potassium chloride (KCl) and oxytocin stimulation. KCl was employed as a receptor-independent positive control to induce smooth muscle contraction by bypassing G-protein-coupled receptors and directly activating voltage-gated calcium channels. Exogenous addition of oxytocin was used to evaluate physiologically relevant uterotonic agonist-mediated myometrial contractility.

At day 3, microtissues failed to generate robust force transients or appreciable increases above baseline force upon stimulation with either 30 mM KCl or 10 µM oxytocin. Representative force traces for KCl (teal) and oxytocin (violet) stimulation are shown in **Figures 4A** and **4B**, respectively, where dotted lines denote day 3 responses. Quantified contraction for KCl and oxytocin is presented in **Figure 4C**, as integrated area under the curve values. Peak contractility is also quantified in **Figure 4D**, as the maximal magnitude of contractile force above baseline.

**Figure 4.**
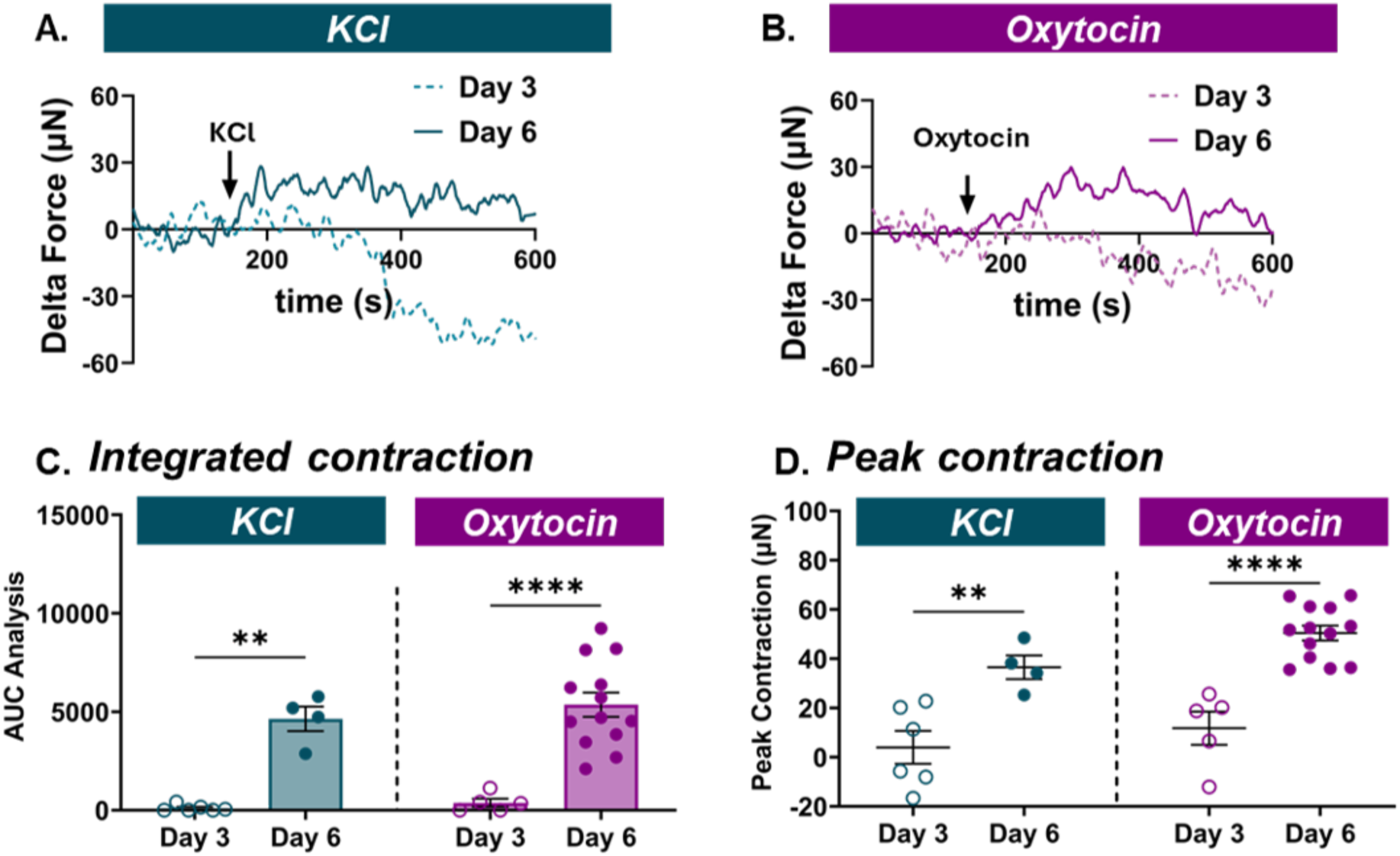
Functional Physiological Response to Exogenous Stimuli of Potassium chloride (KCl) and Oxytocin. Engineered myometrial microtissues were mounted in a force transducer-fitted organ bath and contractile force was measured in response to exogenous addition of potassium chloride (KCl) or oxytocin. Stimulus addition is indicated by the arrow, following establishment of a stable force baseline (arbitrarily set to zero). (A) Representative force traces of microtissue contractile response to 30 mM KCl at day 3 (dotted teal line) and day 6 (solid teal line). (B) Representative force traces of microtissue contractile response to 10 µM oxytocin at day 3 (dotted purple line) and day 6 (solid purple line). (C) Quantification of contractile output as area under the curve (AUC) for KCl and oxytocin stimulation at days 3 and 6. Data are presented as mean ± SEM. ****p<0.0001, **p<0.01; 2-way ANOVA; n ≥ 4. (D) Quantification of mean peak contraction force (µN) following KCl and oxytocin stimulation at days 3 and 6. Data are presented as mean ± SEM. ****p<0.0001, **p<0.01; 2-way ANOVA; n ≥ 4.

By day 6, microtissues displayed progressively stronger and more coordinated contractile responses to both stimuli (compare Day 3 values to Day 6 values in **Figure 4C**). In response to KCl, day 6 microtissues generated a mean peak contraction of 36.53 ± 4.82 µN, representing a 9.16-fold increase relative to day 3 microtissues (**p<0.01, n≥4; 2-way ANOVA; **Figure 4D**). In response to oxytocin, day 6 microtissues generated a mean peak contraction of 50.42 ± 3.03 µN, representing a 4.26-fold increase in peak force relative to day 3 microtissues (****p<0.0001, n≥4; 2-way ANOVA; **Figure 4D**).

### 2.5 Oxytocin induces dose-dependent contractile responses in Engineered Myometrial Microtissues

To determine whether mature myometrial microtissues responded to graded oxytocin dosages, day 6 microtissues were exposed to increasing concentrations of oxytocin (1, 10, and 30 µM) and contractile force was measured (**Figure 5**).

**Figure 5.**
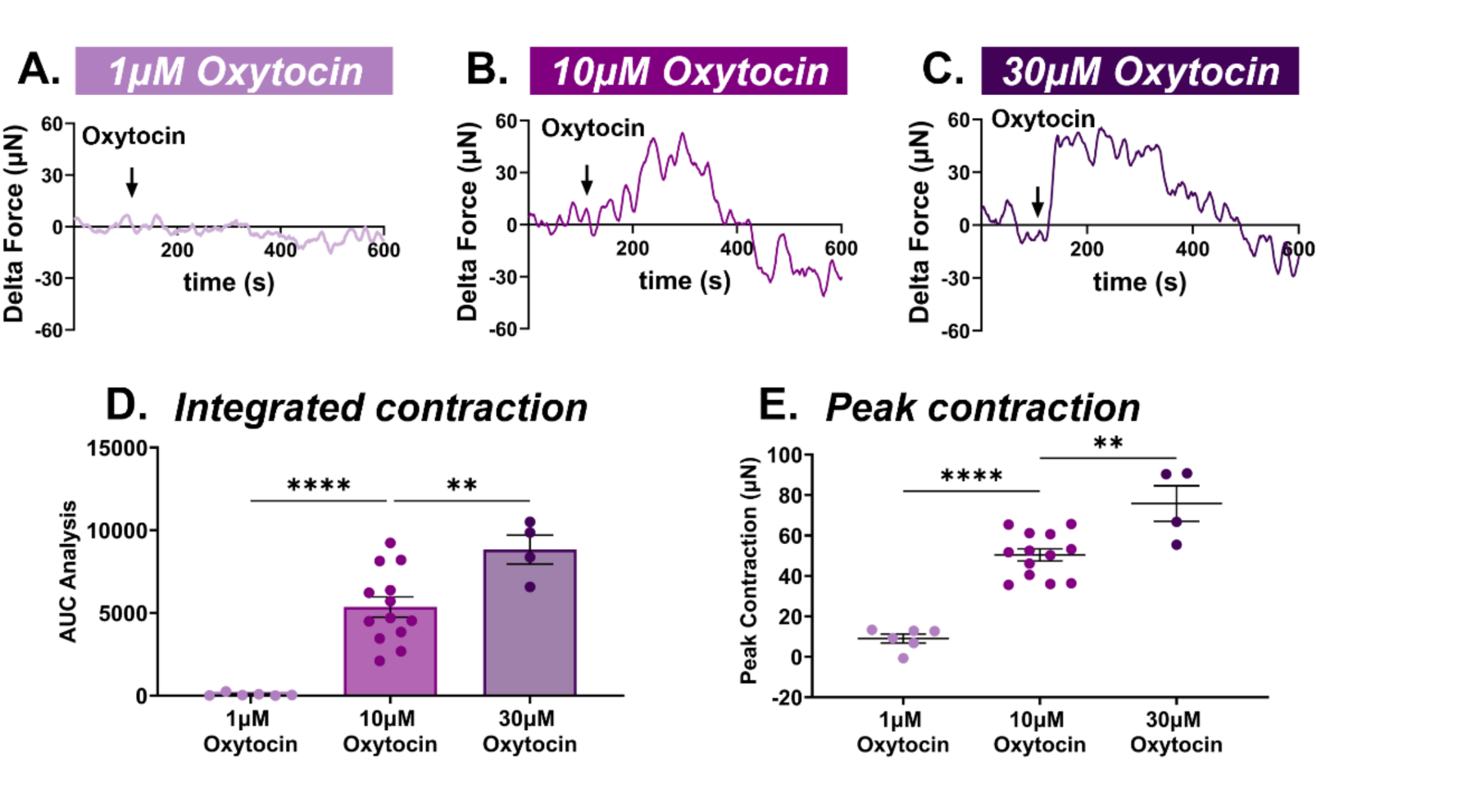
Dose-Dependent Contractile Response to Oxytocin. Representative force traces showing contractile responses of day 6 engineered myometrial microtissues to (A) 1 µM, (B) 10 µM and (C) 30 µM oxytocin. Stimulus addition is indicated by the arrow following establishment of a stable force baseline. (D) Quantification of contractile output as AUC across oxytocin doses (1, 10, and 30 µM). Data are presented as mean ± SEM; n ≥ 4. ****p<0.0001, **p<0.01; one-way ANOVA with Tukey’s multiple comparisons test. (E) Quantification of mean peak contraction force (µN) across oxytocin doses. Data are presented as mean ± SEM; n ≥ 4. ****p<0.0001, **p<0.01; one-way ANOVA with Tukey’s multiple comparisons test.

Representative force traces for 1 µM, 10 µM, and 30 µM oxytocin are shown in **Figures 5A**, **5B**, and **5C**, respectively. At 1 µM oxytocin, microtissues generated a mean maximum peak force of 11.82 ± 6.74 µN. Increasing the concentration to 10 µM resulted in a 73.52-fold increase in total contractile output (quantified as area under the curve in **Figure 5D; \*\*\*\***p<0.0001, n≥4 one-way ANOVA). Corresponding peak contraction with 10µM oxytocin was 5.56-fold higher relative to 1 µM oxytocin (****p<0.0001, n≥4; one-way ANOVA with Tukey’s post-hoc; **Figure 5E**). At the highest tested concentration of 30 µM, microtissues generated a mean peak contraction of 75.85 ± 8.69 µN, corresponding to a 1.50-fold increase in peak contraction relative to the 10 µM condition (**p<0.01, n≥4; one-way ANOVA; **Figure 5E**)

### 2.6 L-Type Calcium Channel Blockade Attenuates KCl- and Oxytocin-Induced Contractility in Engineered Myometrial Microtissues

To determine the role of L-type voltage-gated calcium channel-mediated calcium influx in myometrial microtissue contractility, mature day 6 microtissues were pretreated with the L-type calcium channel blocker nifedipine (10 µM) prior to stimulation with either 30 mM KCl or 10 µM oxytocin (**Figure 6**).

**Figure 6.**
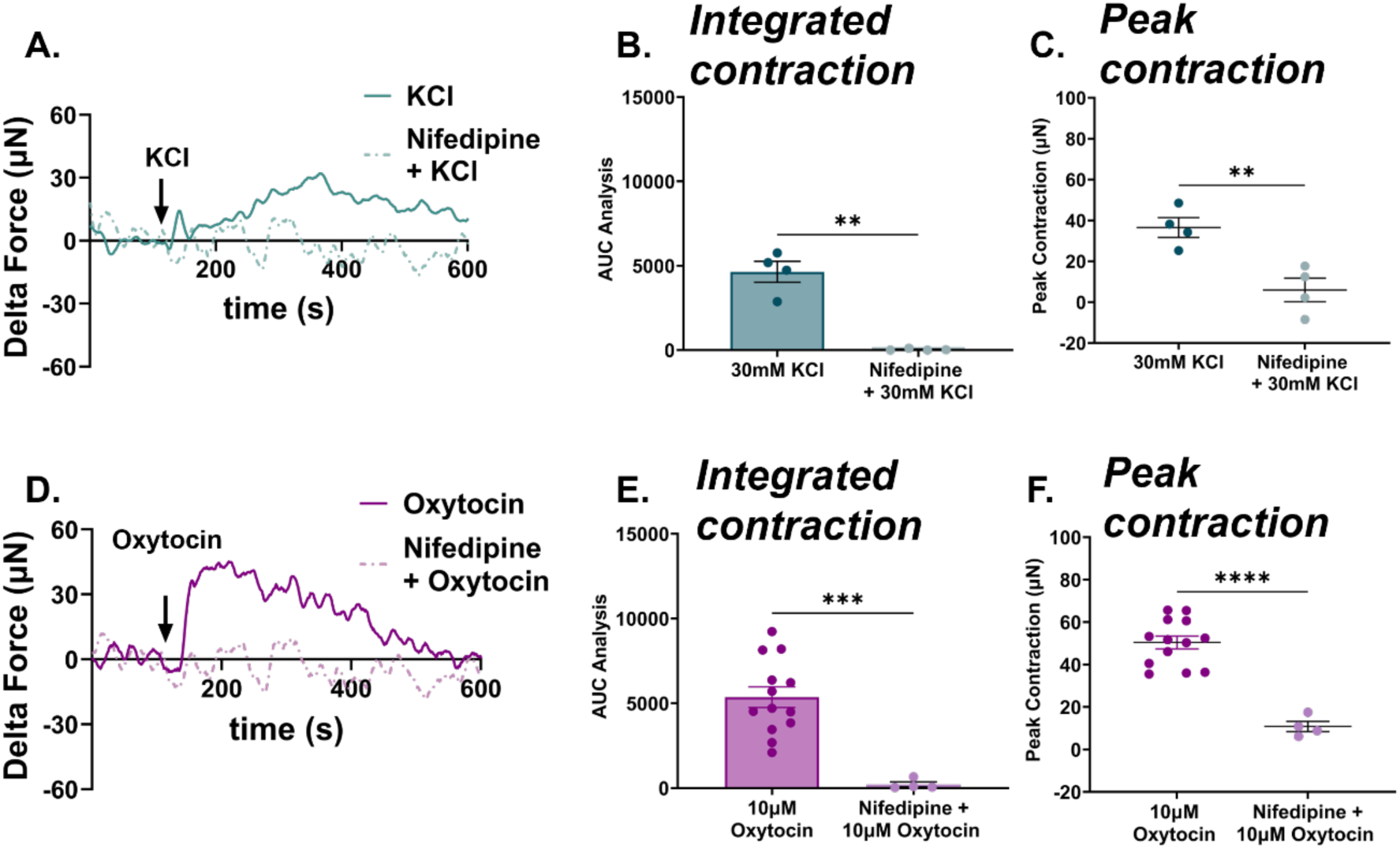
Calcium Dependence of KCl- and Oxytocin-Induced Contractile Responses. Contractile responses of day 6 engineered myometrial microtissues were assessed following pretreatment with the L-type voltage-gated calcium channel blocker nifedipine (10 µM) to evaluate the calcium dependence of KCl- and oxytocin-induced force generation. Stimulus addition is indicated by the arrow following the establishment of a stable force baseline. (A) Representative force traces showing contractile responses to 30 mM KCl alone (solid teal line) or following nifedipine pretreatment prior to KCl stimulation (dashed line). (B) Quantification of contractile output as AUC for KCl versus nifedipine + KCl conditions. Data are presented as mean ± SEM; n = 4. **p<0.01; unpaired t-test. (C) Quantification of mean peak contraction force (µN) for KCl versus nifedipine + KCl conditions. Data are presented as mean ± SEM; n = 4. **p<0.01; unpaired t-test. (D) Representative force traces showing contractile responses to 10 µM oxytocin alone (solid purple line) or following nifedipine pretreatment (dashed line). (E) Quantification of contractile output as AUC for oxytocin versus nifedipine + oxytocin conditions. Data are presented as mean ± SEM; n ≥ 4. ****p<0.0001; unpaired t-test. (F) Quantification of mean peak contraction force (µN) for oxytocin versus nifedipine + oxytocin conditions. Data are presented as mean ± SEM; n ≥ 4. ****p<0.0001; unpaired t-test.

Representative force traces for KCl conditions are shown in **Figure 6A**, with associated area under the curve quantification in **Figure 6B** and peak contractions in **Figure 6C**. Nifedipine pretreatment significantly attenuated KCl-induced contractility, quantified as 99.23% reduction in total contractility (**p<0.01, unpaired t-test; **Figure 6B**), and an 83.75% reduction in peak contractile magnitude (**p<0.01, n≥4; unpaired t-test; **Figure 6C**).

Representative force traces for oxytocin-induced contractility are shown in **Figure 6D**, with or without nifedipine pre-treatment. Nifedipine pretreatment similarly attenuated oxytocin-induced contractility, with a significant 96.1% reduction in total contractility (***p<0.001, n>4; unpaired t-test; **Figure 6E**). Nifedipine pre-treatment resulted in a meagre 10.86 ± 2.41 µN contraction in response to oxytocin, significantly lower than control microtissues stimulated with oxytocin alone and receiving no nifedipine pretreatment (****p<0.0001, n≥4; unpaired t-test; **Figure 6F**).

### 2.7 Oxytocin-Induced Contractility Requires Oxytocin Receptor Signaling

To confirm the specificity of oxytocin-mediated contractility, engineered myometrial microtissues were fabricated using oxytocin receptor knockout (OXTR-KO) hTERT-HM cells (**Supplementary Figures 1 and 2**). Representative force traces for KCl and oxytocin stimulation of OXTR-KO microtissues are shown in **Figures 7A** and 7B.

**Figure 7.**
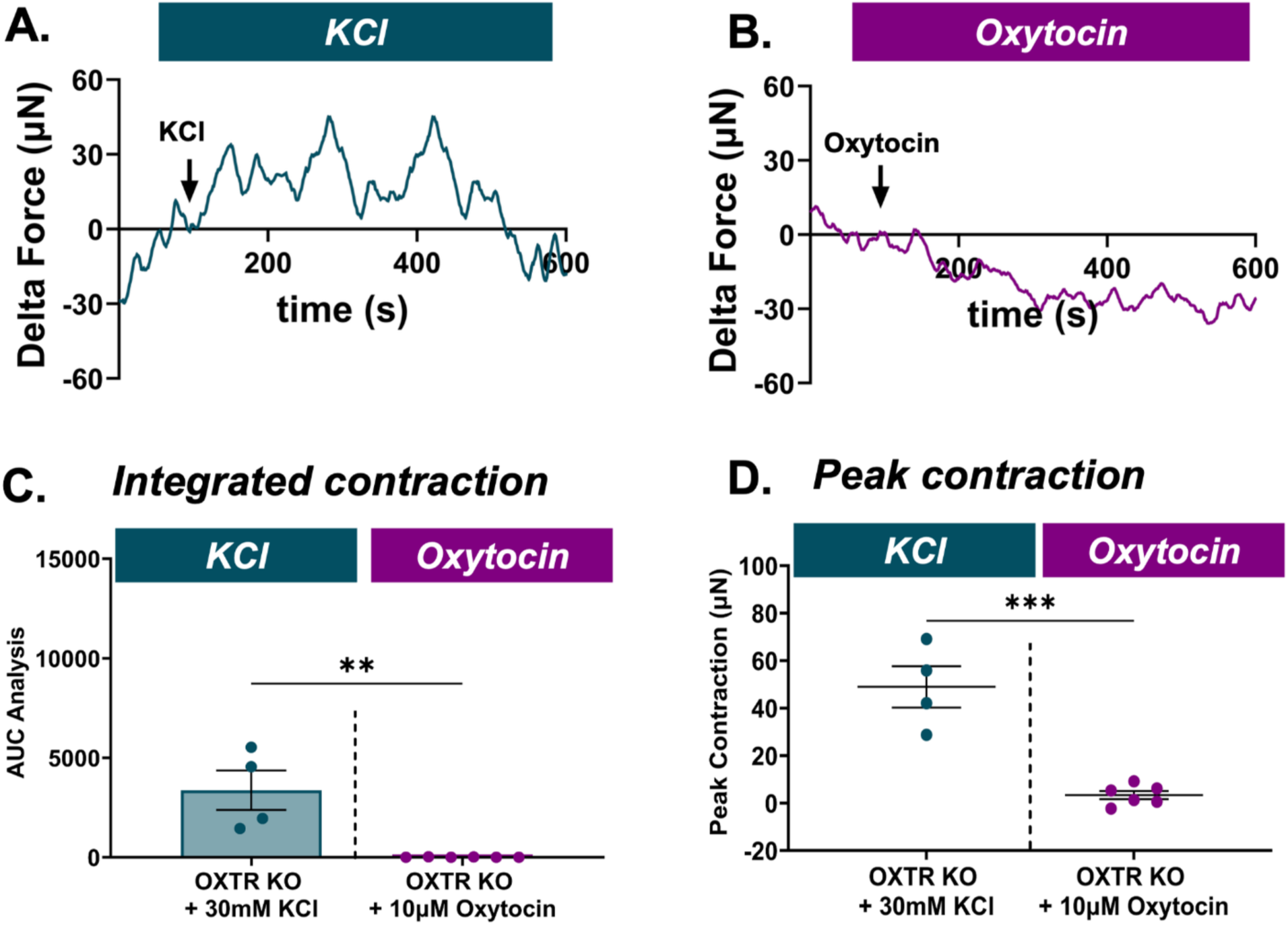
Oxytocin Receptor Specificity of Contractile Responses in Engineered Myometrial Microtissues. (A) Representative force traces of contractile responses in OXTR-KO microtissues stimulated with 30 mM KCl. Stimulus addition is indicated by the arrow following establishment of a stable force baseline. (B) Representative force traces of contractile responses in OXTR-KO microtissues stimulated with 10μM oxytocin. (C) Quantification of contractile output as AUC for OXTR-KO microtissues stimulated with KCl or oxytocin. Data are presented as mean ± SEM; n ≥ 4. *p<0.05; unpaired t-test. (D) Quantification of mean peak contraction force (µN) for OXTR-KO microtissues stimulated with KCl or oxytocin. Data are presented as mean ± SEM; n ≥ 4. *p<0.05; unpaired t-test.

To first confirm that myometrial contractile machinery remained intact in the absence of OXTR, OXTR-KO microtissues were stimulated with 30 mM KCl. OXTR-KO microtissues generated a robust peak contraction of 48.98 ± 8.70 µN in response to KCl stimulation, confirming preserved electromechanical coupling (quantified as total contractility in **Figure 7C**, and peak contractile magnitude in **Figure 7D**).

OXTR-KO microtissues subsequently challenged with 10 µM oxytocin generated a meagre peak contraction of 3.40 ± 1.72 µN, representing a 93.05% reduction relative to KCl-stimulated OXTR-KO microtissues (***p<0.001, n≥4; unpaired t-test; **Figure 7D**). When compared directly to wild-type day 6 microtissues stimulated with 10 µM oxytocin, OXTR-KO microtissues exhibited a significant 93.25% reduction in mean peak contraction (compare contractility of wild-type engineered microtissues in **Figure 4**).

### 2.8 Repeated Oxytocin Exposure Results in Contractile Desensitization with Partial Recovery

To investigate the effects of prolonged oxytocin exposure on contractile responsiveness, mature day 6 microtissues were subjected to continuous oxytocin pre-incubation followed by a defined washout and recovery period, as depicted schematically in **Figure 8A**. Representative force traces for the desensitized and recovery conditions are shown in **Figure 8B**.

**Figure 8.**
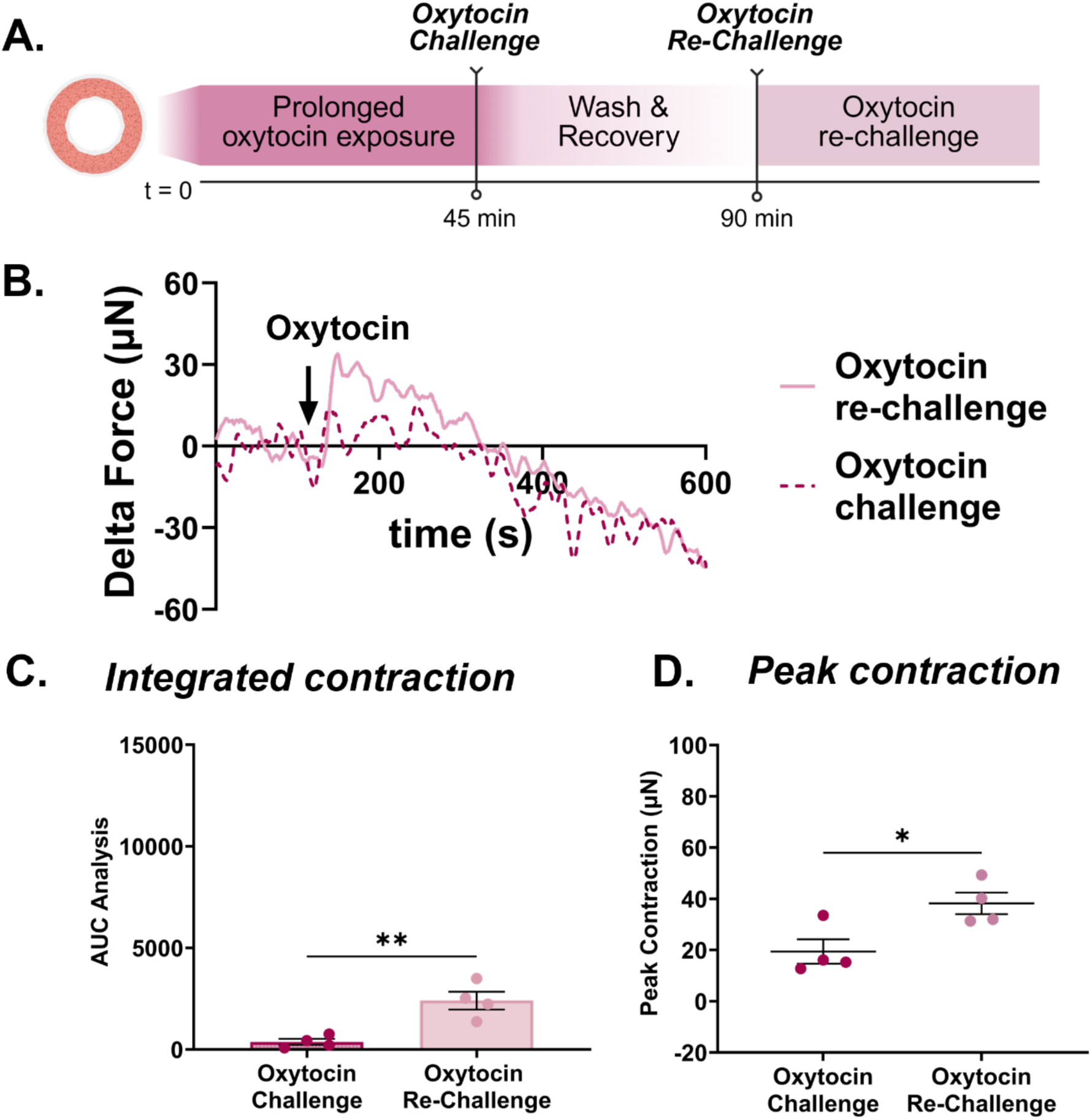
Oxytocin-Induced Contractile Desensitization and Recovery. (A) Schematic of the experimental protocol depicting sequential phases of oxytocin pre-incubation, washout, and recovery. Microtissues were continuously incubated with 10 µM oxytocin prior to stimulation (oxytocin incubation), followed by a 45-minute washout period and subsequent restimulation with 10 µM oxytocin (recovery). (B) Representative force traces of contractile responses during the oxytocin incubation (dotted pink line) and recovery (solid pink line) conditions. Stimulus addition is indicated by the arrow following establishment of a stable force baseline. (C) Quantification of contractile output as AUC for oxytocin incubation and recovery conditions. Data are presented as mean ± SEM; n ≥ 4. *p<0.05; unpaired t-test. (D) Quantification of mean peak contraction force (µN) for oxytocin incubation and recovery conditions. Data are presented as mean ± SEM; n ≥ 4. **p<0.01, *p<0.05; unpaired t-test.

Continuous oxytocin pre-incubation produced a marked attenuation of contractile responses upon subsequent stimulation. Desensitized microtissues generated a mean peak contraction of 19.43 ± 4.76 µN, representing a 90.4% reduction in total contractile output and a 61.46% reduction in mean peak contraction relative to naïve day 6 microtissues stimulated with a single dose of 10 µM oxytocin (**Figure 8D**).

Following a 45-minute washout and recovery period, microtissues partially regained contractile responsiveness, generating a mean peak contraction of 38.21 ± 4.20 µN, corresponding to a 1.97-fold increase relative to the desensitized state (*p<0.05, n≥4; unpaired t-test; **Figure 8D**). Nevertheless, mean peak contraction following recovery remained 24.22% below naïve baseline values, indicating incomplete restoration of contractile responsiveness within the observation window.

## 3. DISCUSSION

This study presents a functional three-dimensional human myometrium microtissue model (EMMI) that enables direct, quantitative measurement of isometric contractile force in response to contractile agonists and stimuli. Unlike existing three-dimensional systems that assess contraction indirectly via changes in tissue area, diameter, or collagen gel compaction, this platform provides a direct readout of force generation, which remains the physiological gold standard established by human myometrial tissue strip studies(*10, 14, 24, 29*). By integrating cellular self-assembly and real-time force generation measurements, the engineered myometrial microtissues recapitulate key structural, molecular, and functional features of native human myometrial tissue.

Animal models have provided important mechanistic insights into uterine physiology; however, fundamental species-specific differences in reproductive biology, including the timing and hormonal triggers of labor onset, uterine architecture, and contractile waveform characteristics, substantially limit their translational relevance to human pregnancy and parturition(*30, 31*). The present platform, derived entirely from human myometrial cells, circumvents these limitations and provides a species-appropriate context for interrogating contractile signaling pathways.

The engineered myometrial microtissues (EMMIs) underwent coordinated maturation characterized by progressive matrix compaction, cytoskeletal alignment, and increased expression of smooth muscle contractile and junctional proteins. Rapid hydrogel compaction, most pronounced between day 0 and day 1, confirmed that hTERT-HM cells were mechanically active and capable of remodeling the extracellular matrix from the earliest stages of culture. Whole-mount immunofluorescence imaging provided microscopic insight into the evolving architecture of the microtissues, revealing progressive circumferential alignment of elongated, spindle-shaped myocytes resembling the circular layer of the native myometrium, where circumferentially oriented smooth muscle cells interconnected by gap junctions generate coordinated uterine contractions(*2, 13, 32*). Bulk RNA sequencing of the microtissues provided transcriptomic evidence of their time-dependent maturation. The transcriptome of the day 6 microtissues was distinct from that of day 3 and 1 tissues, matching the change in contractile potential of the microtissues at this time point. Similarly, in native myometrial tissue, the large transcriptomic changes occurring at term are accompanied by a significant shift in contractile potential of the organ(*33*). In native tissue, these transcriptomic changes coordinate a shift from a proliferative to a contractile phenotype in the myometrial smooth muscle cells(*34*). At day 3 and 6, the microtissues demonstrated an upregulation of *ACTA2*, a known smooth muscle cell marker and an enrichment in muscle-related biological pathways(*35*). These shifts highlight transcriptional changes in cellular machinery that are consistent with a time-dependent shift toward a mature, contractile smooth muscle phenotype, mirroring differentiation processes observed in native smooth muscle tissue(*36*).

Furthermore, the scaffold-free architecture of the EMMIs minimizes mechanical mismatch between synthetic matrices and contractile tissue, a factor known to influence force development in muscle systems(*37, 38*). The self-assembling nature of the platform supports more physiologically relevant force generation, as cells remodel and compact their own extracellular matrix rather than being constrained by an exogenous scaffold(*39*). The ring geometry structurally models the circular layer of the myometrium, which is critical for coordinated uterine contraction, and enables repeated, longitudinal functional assessments within a single microtissue, reducing inter-sample variability and tissue consumption.

A major strength of the EMMI platform is its tunability to interrogate specific aspects of myometrial physiology. In this study, we focused on oxytocin-mediated contractile signaling due to its central importance in uterine tone regulation and parturition. Mature microtissues exhibited dose-dependent contractile responses to oxytocin, confirming functional oxytocin receptor coupling to downstream contraction pathways(*40, 41*). The oxytocin concentrations required to elicit robust contractile responses in the engineered microtissues (10 to 30 µM) were higher than those typically reported in *ex vivo* myometrial tissue strip studies, where significant responses are observed at concentrations as low as 1 to 10 nM(*42, 43*), and substantially higher than the low nanomolar oxytocin exposure achieved during clinical labor induction protocols(*44–46*). This shift in dose-response may be due to two converging factors. First, engineered microtissues contain a substantially lower mass of contractile tissue and fewer smooth muscle fascicles than intact *ex vivo* tissue strips, reducing total force-generating capacity per unit of agonist. Second, hTERT-HM cells originated from non-pregnant myometrium and were maintained under non-pregnant culture conditions, which are associated with markedly lower oxytocin receptor expression compared to term pregnant myometrial tissue from which *ex vivo* strips are typically derived(*7, 8*). These findings are consistent with previous reports of elevated oxytocin concentrations required to elicit responses in engineered myometrial tissue systems(*21*) and highlight hormonal preconditioning to upregulate oxytocin receptor expression as an important priority for future development of the platform.

The engineered myometrial microtissue platform also recapitulates key regulatory features of human myometrial contractile physiology. Pharmacological blockade of L-type voltage-gated calcium channels with nifedipine markedly attenuated both KCl- and oxytocin-induced contractility, confirming the central role of calcium influx through L-type channels in excitation-contraction coupling in the engineered microtissues, consistent with established myometrial physiology(*47–49*). In the native myometrium, membrane depolarization triggers calcium influx through L-type voltage-gated calcium channels, which activates myosin light chain kinase, drives actin-myosin cross-bridge cycling, and ultimately generates contractile force, a pathway that is critically dependent on extracellular calcium availability and is the primary target of tocolytic agents such as nifedipine in the clinical management of preterm labor(*47, 48*). The receptor independence of KCl-induced contraction, combined with its near-complete abolition by nifedipine pretreatment, further confirms that force generation in this platform is dependent on voltage-gated calcium influx in addition to receptor-mediated signaling.

A key advantage of the engineered myometrial microtissue platform over traditional *ex vivo* tissue strip preparations is its compatibility with genetic manipulation, enabling mechanistic dissection of specific signaling pathways that is not feasible with primary human tissue strips. To demonstrate this capability, microtissues were fabricated from OXTR-KO hTERT-HM cells, allowing direct interrogation of oxytocin receptor-specific contractile signaling in a three-dimensional force-generating system. Genetic deletion of the oxytocin receptor markedly attenuated oxytocin-induced contractility, while KCl-induced depolarization-driven contractions remained intact, confirming that oxytocin-mediated force generation is driven by specific oxytocin receptor signaling rather than nonspecific calcium influx or matrix-mediated effects. Such receptor-specific genetic validation is inherently impossible in *ex vivo* tissue strips derived from human surgical specimens, highlighting the unique mechanistic interrogation capabilities of this platform.

Repeated oxytocin stimulation resulted in progressive contractile desensitization, with a marked attenuation of force generation following continuous oxytocin pre-incubation, consistent with oxytocin receptor desensitization and signaling exhaustion observed in clinical and *ex vivo* studies(*7, 8, 11*). Partial recovery of contractile responsiveness was observed following a 45-minute washout period, mirroring the incomplete receptor resensitization reported in *ex vivo* human myometrial tissue strip studies following oxytocin withdrawal(*50*). The incomplete restoration of contractile output within the observation window is consistent with the known timescale of receptor resensitization, which proceeds primarily via recycling of internalized receptors to the plasma membrane through a Rab4/Rab5-dependent pathway(*51, 52*). Accordingly, the 45-minute washout period represents a minimum estimate of resensitization capacity; given more time, more complete receptor recycling and contractile recovery would be expected. This partial but measurable recovery mirrors the clinical observation that oxytocin-induced uterine atony and impaired contractility can persist well beyond the cessation of oxytocin infusion during labor(*3–6*). Critically, these dynamic desensitization and recovery behaviors emerge in a simplified, controllable system, enabling mechanistic dissection of oxytocin receptor regulation that is not readily achievable in human tissue strips or *in vivo* models.

The use of non-pregnant human myometrial cells as the cellular basis for this platform represents a deliberate design choice that expands the translational scope of the engineered myometrial microtissue platform beyond term pregnant physiology. Non-pregnant myometrium expresses functional oxytocin receptors that drive junctional zone peristalsis and are implicated in the pathophysiology of endometriosis, fibroids, and implantation failure(*15, 53–55*). Because most existing *ex vivo* strip studies are restricted to term pregnant tissue obtained at cesarean delivery, their findings may not generalize to the broader range of clinical conditions that affect myometrial function across the reproductive lifespan. The use of an immortalized non-pregnant cell source therefore enables controlled interrogation of contractile signaling pathways relevant to pregnancy, postpartum states, uterine fibroids, adenomyosis, and other myometrial pathologies without the constraints of limited surgical specimen availability. Furthermore, the immortalized nature of the hTERT-HM cell line ensures a reproducible, genetically stable cellular substrate that can be consistently expanded and manipulated across laboratories, addressing a key reproducibility limitation of primary cell-based models.

Despite these strengths, several limitations of the engineered myometrial microtissue platform warrant consideration. The microtissues are composed of a single cell type and therefore do not capture the contributions of fibroblasts, immune cells, or endothelial cells present in native myometrial tissue, which may influence contractile responses through paracrine signaling and matrix remodeling. Additionally, the extracellular matrix is limited to type I collagen and does not fully recapitulate the biochemical or mechanical complexity of the native uterine extracellular matrix environment(*15*). The use of non-pregnant hTERT-HM cells, while advantageous for reproducibility and broad translational applicability, does not capture the hormonal milieu of pregnancy, including the functional progesterone withdrawal and associated upregulation of oxytocin receptors that characterize term parturition. Future studies incorporating use of primary myometrial cells, multicellular co-cultures, tunable matrix compositions, and hormonal preconditioning will further enhance the physiological relevance and translational applicability of this platform.

## 4. CONCLUSION

The engineered myometrial microtissues presented here recapitulate key structural, molecular, and functional features of native human myometrial tissue in a reproducible, genetically tractable, and directly quantifiable system. By linking molecular maturation to isometric force generation, and by demonstrating pharmacologically specific contractile responses to oxytocin and KCl, calcium channel dependency, receptor-specific signaling, and clinically relevant oxytocin desensitization, this platform establishes a new framework for investigating uterine contractile biology. Future studies incorporating multicellular co-cultures, hormonal preconditioning, and tunable matrix compositions will further enhance the physiological relevance of this system. Ultimately, the engineered myometrial microtissue platform provides a versatile and mechanistically tractable tool for interrogating myometrial physiology, modeling uterine pathologies, and evaluating novel therapeutic strategies for disorders ranging from preterm labor and uterine atony to fibroids and adenomyosis.

## 5. MATERIALS AND METHODS

### 5.1 Materials and reagents

All cell culture media and supplements were purchased from Thermo Fisher Scientific (Waltham, MA, USA) unless otherwise specified.

Oxytocin was purchased from Tocris Bioscience (Cat# 1910). Nifedipine was purchased from Sigma-Aldrich (Cat# N7634-1G). Type I rat tail collagen was purchased from ibidi (Madison, WI). Polydimethylsiloxane (PDMS; Sylgard 184 Silicone Elastomer Kit) was purchased from Dow Corning (Midland, MI). Paraformaldehyde (4%) was purchased from Santa Cruz Biotechnology (Cat# sc-281692). Bovine serum albumin (BSA) was purchased from Fisher Bioreagents (Cat# BP9706-100). Anti-connexin-43 primary antibody was purchased from Sigma-Aldrich (Cat# C6219, RRID:AB_476857). Anti-smoothelin primary antibody was purchased from Novus Biologicals (Cat# NBP2-37931, RRID:AB_3297445). Anti-OXTR primary antibody was purchased from Santa Cruz Biotechnology (Cat# SC-515809, RRID: AB_2891171). Revert™ 700 Total Protein Stain was purchased from LI-COR Biosciences (P/N 926-11021, Lincoln, NE). The RNeasy Mini Kit was purchased from Qiagen (Hilden, Germany).

### 5.2 Cell lines and culture

#### Culture Of Immortalized Human Myometrium Smooth Muscle Cells (hTERT-HM)

Immortalized human myometrial smooth muscle cells (hTERT-HM) were maintained in phenol red-free DMEM/F12 supplemented with 10% fetal bovine serum (FBS) and 25 µg/mL gentamicin(*16*). Cultures were maintained at 37°C in a humidified atmosphere of 5% CO₂ and passaged below 80% confluency using 0.25% trypsin-EDTA. All experiments were performed using cells under passage 17, to ensure phenotypic consistency.

#### Developing stable knockout cell lines

The OXTR KO hTERT-HM cell line was generated using clustered regularly interspaced short palindromic repeat (CRISPR) Cas9 nuclease as described elsewhere(*56*). Briefly, oligonucleotide encoding the guide RNA (*gacatcaccttccgcttcta*) were cloned into linearized GeneArt CRISPR Nuclease/CD4 Enrichment vector, which also encoded Cas9 and CD4 genes (Cat# A21175, Life Technologies). Sequence-confirmed plasmid was transfected into hTERT-HM cells using Lipofectamine 3000 Transfection Reagent (Cat# L3000001, ThermoFisher Scientific). At 24 hours post transfection, Dynabeads CD4 isolation kit (Cat# 11331D, Life Technologies) was used to enrich for cells expressing surface CD4. Individual single cell-derived clones were selected, propagated, and analyzed for oxytocin response using the Brillian Calcium Flex assay (Cat# 1000, IONBiosciences). Clones with a negative response to oxytocin stimulation were then also analyzed for expression of OXTR protein by immunoblot (**Supplementary Figure 1**).

### 5.3 Fabrication of Engineered Myometrial Microtissues (EMMIs)

EMMIs were fabricated through a previously established collagen hydrogel deposition technique to enable self-assembly of smooth muscle cells within cell-laden hydrogels(*57*). Briefly, 35 mm dishes were prepared by coating with a thin layer of polydimethylsiloxane (PDMS; Sylgard^TM^ 184 silicone elastomer kit, Dow Inc., Midland, MI). A central silicone post (3 mm diameter) was embedded within the PDMS layer to define and maintain a hollow luminal space. Following curing for 48 hours, dishes presented a non-adherent surface with a defined central luminal post geometry.

hTERT-HM cells were uniformly suspended within a collagen hydrogel (1mg/mL) at a density of 1.67 × 10⁶ cells/mL. 600 μL of cell-laden gel was cast around the central post and allowed to polymerize at 37°C for 30 minutes. Following gelation, 2 mL of hTERT-HM culture medium was added to each dish. EMMIs were maintained at 37°C to promote cellular self-assembly, alignment, and gel compaction. Microtissue formation was monitored visually over 6 days, and compaction of the gel area was quantified as described in the following section.

### 5.4 Compaction analysis

To monitor EMMI compaction over time, photographs of each construct were captured daily from day 0 to day 6 using a consistent overhead setup with an iPhone camera. Photos were captured from a standardized fixed distance and angle above the culture dish, under consistent ambient lighting conditions, to ensure reproducibility across all timepoints. Images were imported into Fiji software(*58*) and calibrated for scale using the known inner diameter of the 35 mm culture dish as a spatial reference, with the scale bar set accordingly prior to analysis. Briefly, the outer boundary of the collagen gel was manually traced using the freehand selection tool, and the area occupied by the central silicone post was subsequently subtracted to isolate the net gel area. This process was repeated for each daily image across all constructs. Compaction was normalized to the initial gel area measured at day 0 for each individual construct and reported as percent change over time.

### 5.5 Characterization of structural maturity of EMMIs using wholemount immunofluorescence

EMMIs were fixed at days 1, 3, and 6 in 4% paraformaldehyde (Santa Cruz Biotechnology Cat# sc-281692) for 30 minutes at room temperature (RT) and subsequently permeabilized in 0.1% Triton-X-100 (Thermo Fisher Scientific Cat# A16046.AE) in 1x phosphate-buffered saline (PBS) overnight at 4°C. Microtissues were blocked in 5% bovine serum albumin (BSA; Fisher Bioreagents Cat# BP9706-100) overnight at 4°C.

To assess gap junction formation, smooth muscle cell maturity, and actin organization, EMMIs were stained for connexin-43 (Cx43), smoothelin, and filamentous actin (phalloidin), respectively. For Cx43 and smoothelin, microtissues were incubated in primary rabbit antibodies against connexin-43 (1:200; Sigma-Aldrich Cat# C6219, RRID:AB_476857) or smoothelin (1:100; Novus Cat# NBP2-37931, RRID:AB_3297445) overnight at 4°C and subsequently incubated in anti-rabbit secondary antibody (1:200; Thermo Fisher Scientific Cat# SA5-10041, RRID:AB_2556621) overnight at 4°C. For actin alignment, EMMIs were incubated overnight at 4°C in phalloidin labeling probes (1:150; Thermo Fisher Scientific Cat# A34054). All microtissues were counterstained with DAPI for 20 minutes at RT for nuclear visualization.

After staining, EMMIs were imaged with a fluorescent confocal microscope (Nikon ECLIPSE Ti2 inverted microscope, RRID:SCR_021068). Three microtissues per timepoint were stained and imaged, with three regions of interest (ROIs) captured per microtissue to account for intra-construct variability.

### 5.6 Quantification of actin alignment

To quantify cytoskeletal alignment over time, coherency of phalloidin-stained actin fibers was measured using the OrientationJ plugin (http://bigwww.epfl.ch/demo/orientation/) in Fiji(*58, 59*). Images were first converted to 8-bit, and OrientationJ Analysis was applied using default parameters to generate orientation colormaps, which were used to visually verify accurate fiber detection. Fiber coherency was then quantified using the OrientationJ Dominant Direction function, where a coherency value of 0 indicates fully isotropic, randomly oriented fibers, and a value of 1 indicates highly aligned, unidirectional fibers. Coherency values across the three ROIs per microtissue were averaged, and these averaged values were used for statistical analysis.

### 5.7 Quantification of structural and contractile maturity of cell lines and EMMIs using western blotting

EMMIs were washed with phosphate-buffered saline (PBS). Tissues were sonicated (CL-188 immersion probe QSonica, Newton, CT) on ice in 200µL of lysis buffer (150mM NaCl, 50mM Tris HCl [pH 8.0], 1mM EDTA, 0.5% Nonidet P-40, 2% Glycerol, 1% Halt Protease & Phosphatase Inhibitor Cocktail [Thermo Fisher Scientific, Waltham, MA]). Lysates rested on ice for 10 minutes before being centrifuged at 9000g for 10 min at 4°C. Supernatants were stored at −20°C. Protein concentration was determined using the Pierce BCA Protein Assay Kit (Thermo Fisher Scientific, Waltham, MA). Lysates were separated via SDS-PAGE in 4%-12% Bolt Bis-Tris gels (Thermo Fisher Scientific, Waltham, MA). Protein was transferred to PVDF membranes, which were then blocked in Blocker FL Fluorescent Blocking Buffer (Thermo Fisher Scientific, Waltham, MA) for 1 hour. Membranes were incubated overnight at 4°C with primary antibodies and then incubated with appropriate secondary antibodies at RT for 1 hour. Membranes were stained with antibodies against: smoothelin (1:1000, Novus Biologicals, NBP2-37931, RRID:AB_3297445), connexin-43 (1:5000, Sigma C6219, RRID:AB_476857) or GAPDH (1:10,000, Protein Tech 60004-1-Ig, RRID:AB_2107436), followed by anti-mouse IgG (H + L) (Jackson Immunoresearch Labs Cat #715-065-150, RRID: AB_2307438) or Donkey anti-Rabbit IgG (H + L) Alexa Fluor Plus 800 (Invitrogen Cat #A32808, RRID: AB_2762837).

Signal was detected with ECL Western Blot Substrate (Thermo Fisher Scientific, Waltham, MA) when appropriate. Membranes were scanned using ChemiDoc MP (Bio-Rad Laboratories, Hercules, CA)) and quantitated in Image Lab (Bio-Rad Laboratories). Values for smoothelin and connexin-43 were normalized to GAPDH.

Similarly, lysates were collected from multiple passages of WT and OXTR KO hTERT-HM cells and immunoblotting was performed. Before blocking, total protein was detected using REVERT Kit (Li-COR) following manufacturer instructions. After REVERT staining membrane was washed with DI water and incubated with appropriate primary and secondary antibodies as described above. Values for smoothelin, connexin-43, and OXTR were normalized to total protein.

### 5.8 Functional contractility assessment using a force-transducer based organ bath

Contractile function of EMMIs was assessed using a custom-built organ bath system, as previously described(*57*). Each microtissue was mounted by securing one end to a fixed reference post and the other to the arm of a force transducer (FT20, Harvard Apparatus). The organ bath was filled with DMEM buffered with 1 M HEPES and maintained at 37°C throughout the experiment. A 30% passive stretch was applied via micromanipulator and maintained for the duration of the recording to measure isometric force. Microtissues were equilibrated under these conditions for 60-90 minutes prior to pharmacological stimulation. Baseline force following equilibration was set to zero to allow direct comparison of pharmacologically induced contractile responses.

To characterize smooth muscle contractile function, EMMIs were treated with potassium chloride (KCl; 30 mM) to induce membrane depolarization-driven contraction, or oxytocin (1-30μM) to assess uterotonic agonist-mediated contractility. Pharmacological agents were administered directly into the bath, and contractile responses were recorded continuously. Force traces were analyzed for peak amplitude and area under the curve to quantify contractile magnitude and duration, respectively. Each condition was assessed across a minimum of four biological replicates (n ≥ 4).

### 5.9 RNA Sequencing and analysis

Total RNA was extracted from EMMIs at days 1, 3, and 6 using the RNeasy Mini Kit (Qiagen, Hilden, Germany) according to the manufacturer’s instructions. RNA concentration and purity were assessed using a NanoDrop 2000 spectrophotometer (Thermo Fisher Scientific, Waltham, MA). Extracted RNA was subsequently submitted for RNA sequencing to the Washington University Genome Technology Access Center at the McDonnell Genome Institute for library preparation and sequencing. Total RNA integrity was determined on an Agilent Bioanalyzer, and all samples had an RNA integrity score above 8.0. Libraries were generated with 1µg of total RNA. Ribosomal RNA was removed via poly-A-selection with Oligo-dT beads. mRNA was fragmented with reverse transcriptase buffer and heating to 94°C for 8 minutes. SuperScript III RT enzyme (Life Technologies) and random hexamers were used per manufacturer instructions to reverse transcribe mRNA into cDNA. A second reaction was performed to yield ds-cDNA. Blunt ended cDNA had an A base added to the 3’ end and had Illumina sequencing adapters ligated to the ends. Ligated fragments were amplified for 12-15 cycles using primers incorporating dual index tags. Fragments were sequenced on an Illumina NovaSeq X Plus using paired end reads extending 150 bases. Basecalls and demultiplexing were performed with Illumina’s DRAGEN BCLconvert version 4.3.13. RNA-seq reads were aligned to the Ensembl GRCh38.113 primary assembly with STAR version 2.7.11b. Gene counts were derived from the number of uniquely aligned unambiguous fragments by Subread:featureCount version 2.0.8. The RNAseq data (DOI pending) were submitted to the NCBI Gene Expression Omnibus database ((*60*)).

DESeq2 was used to identify differentially expressed genes (DEGs) from gene counts(*61*). Genes with an adjusted p-value (padj) < 0.01 and a log_2_fold change > 0.58 were considered significant. Gene ontology analysis was performed on the DEGs using the clusterProfiler R package(*62*).

### 5.10 Statistical Analysis

All statistical analyses were performed using GraphPad Prism 11 (GraphPad Software, San Diego, CA) unless stated otherwise. Data are presented as mean ± standard error of the mean (SEM). Data for each instance was generated from at least three biological replicates of engineered myometrial microtissues where indicated. ANOVA-based hypothesis testing with appropriate post-hoc tests were used to compare between groups, with statistical significance reported at each instance.

## 7. ACKNOWLEDGEMENTS

## 7.3 Author Contributions

Karla I. Ortega Sandoval: Methodology, investigation, formal analysis, data curation, visualization, writing – original draft, writing – review & editing

Ritu M. Dave: Methodology, investigation, formal analysis, data curation, writing – review & editing

Cailin R Gonyea: Investigation, formal analysis, data curation, writing – review & editing Kaci Mitchum: Investigation

Alana Aristimuno Millan: Investigation, Visualization Shriya Suryakumar: Investigation, Visualization

Antonina I. Frolova: Conceptualization, methodology, funding acquisition, project administration, resources, supervision, validation, visualization, writing – review & editing

Shreya A. Raghavan: Conceptualization, methodology, funding acquisition, project administration, resources, supervision, validation, visualization, writing – original draft, writing – review & editing

## 7.4 Data Availability

Raw and processed data supporting the findings of this study is being deposited in the Texas Data Repository, with a DOI ahead of publication. RNA Sequencing data will be deposited in the GEO Database, with a DOI made publicly available.

## Supplementary Information

Supplementary file: Additional information describing Oxytocin receptor knock out cell lines used in the main manuscript.

**Supplementary Figure 1.**
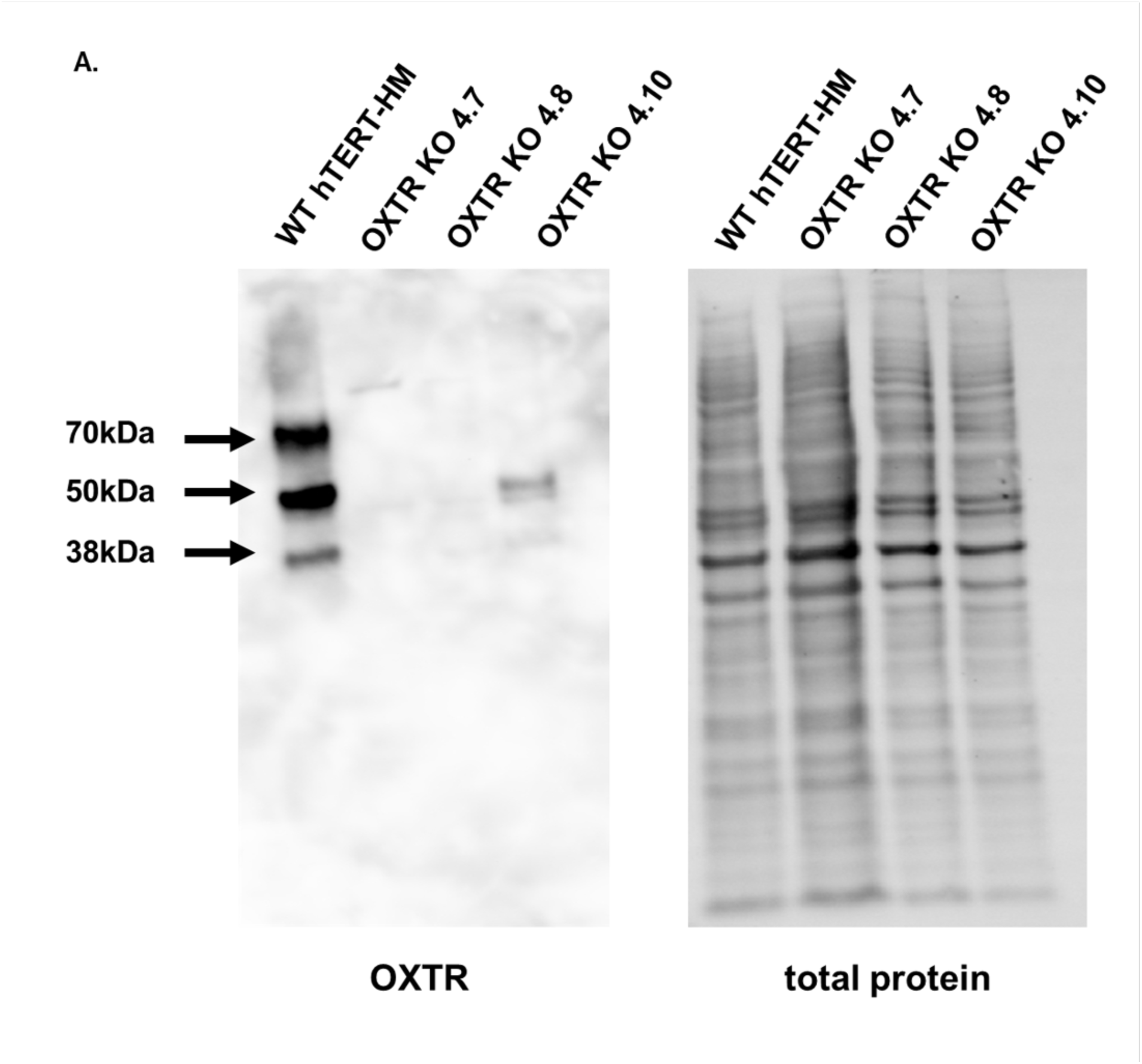
*Validation of OXTR Protein Loss in CRISPR-Generated hTERT-HM Knockout Clones* (A)Representative western blot confirming loss of oxytocin receptor (OXTR) protein expression in OXTR KO hTERT-HM clone compared to wild-type (WT) hTERT-HM cells (left panel), OXTR protein is present in WT hTERT-HM cells and absent in clones OXTR KO 4.7, 4.8 and 4.10. Total protein staining confirms equal protein loading across all lanes (right panel)

**Supplementary Figure 2.**
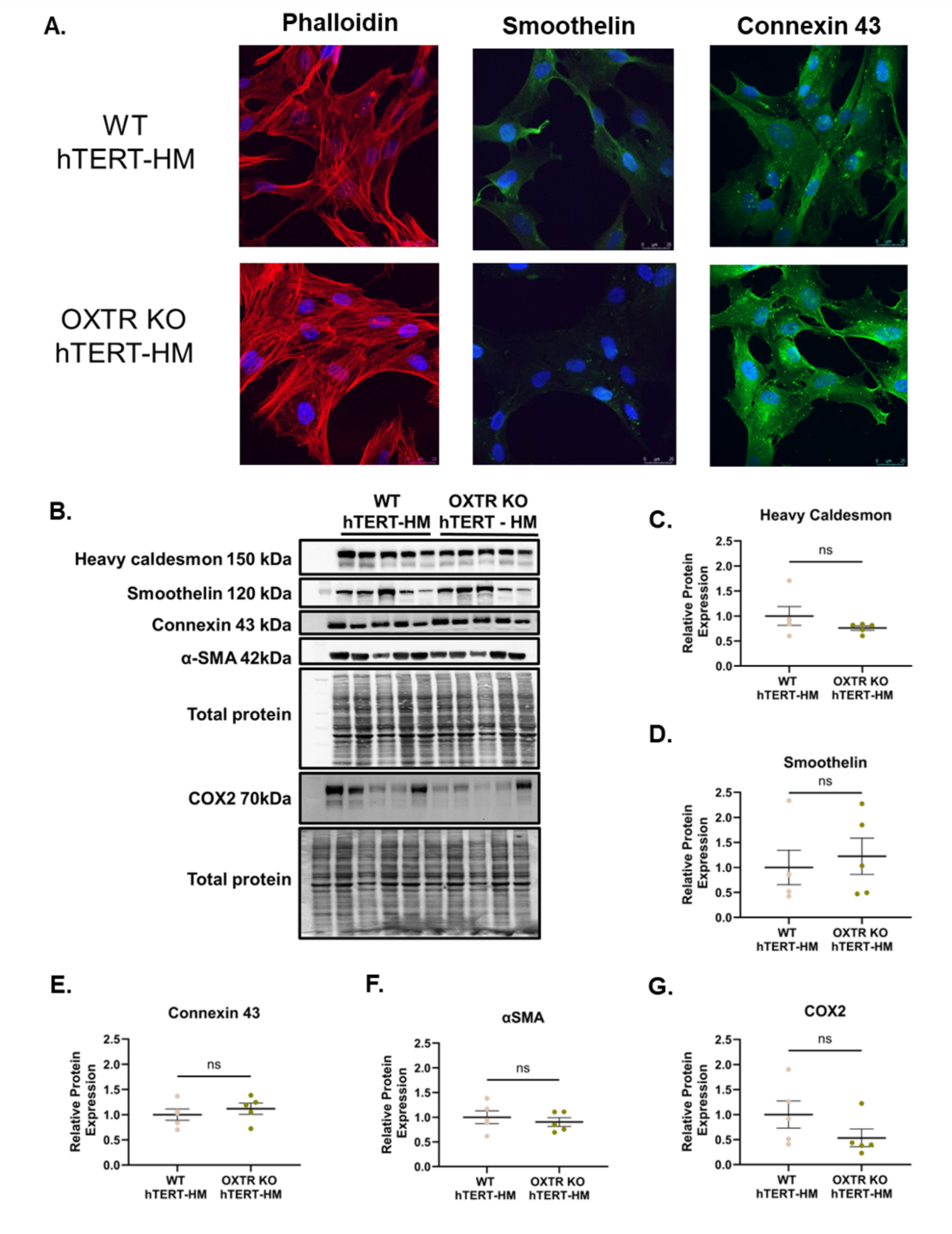
Characterization of OXTR-KO hTERT-HM Cells. (A) Representative whole-mount immunofluorescence images of wild-type (WT) and oxytocin receptor knockout (OXTR-KO) hTERT-HM cells stained for F-actin (phalloidin, red), smoothelin (green), and connexin-43 (green). Nuclei are counterstained with DAPI (blue). Scale bar = 25 µm. (B) Representative western blots for heavy caldesmon (150 kDa), smoothelin (120 kDa), connexin-43 (43 kDa), α-smooth muscle actin (α-SMA; 42 kDa), and COX2 (70 kDa) in WT and OXTR-KO hTERT-HM cells. Total protein is shown as loading control. (C) Quantification of heavy caldesmon relative protein expression normalized to total protein in WT and OXTR-KO hTERT-HM cells. Data are presented as mean ± SEM; n = 3. ns, not significant; unpaired t-test (D) Quantification of smoothelin relative protein expression normalized to total protein in WT and OXTR-KO hTERT-HM cells. Data are presented as mean ± SEM; n = 3. ns, not significant; unpaired t-test. (E) Quantification of connexin-43 relative protein expression normalized to total protein in WT and OXTR-KO hTERT-HM cells. Data are presented as mean ± SEM; n = 3. ns, not significant; unpaired t-test. (F) Quantification of α-SMA relative protein expression normalized to total protein in WT and OXTR-KO hTERT-HM cells. Data are presented as mean ± SEM; n = 3. ns, not significant; unpaired t-test. (G) Quantification of COX2 relative protein expression normalized to total protein in WT and OXTR-KO hTERT-HM cells. Data are presented as mean ± SEM; n = 3. ns, not significant; unpaired t-test.

**Supplementary Figure 3.**
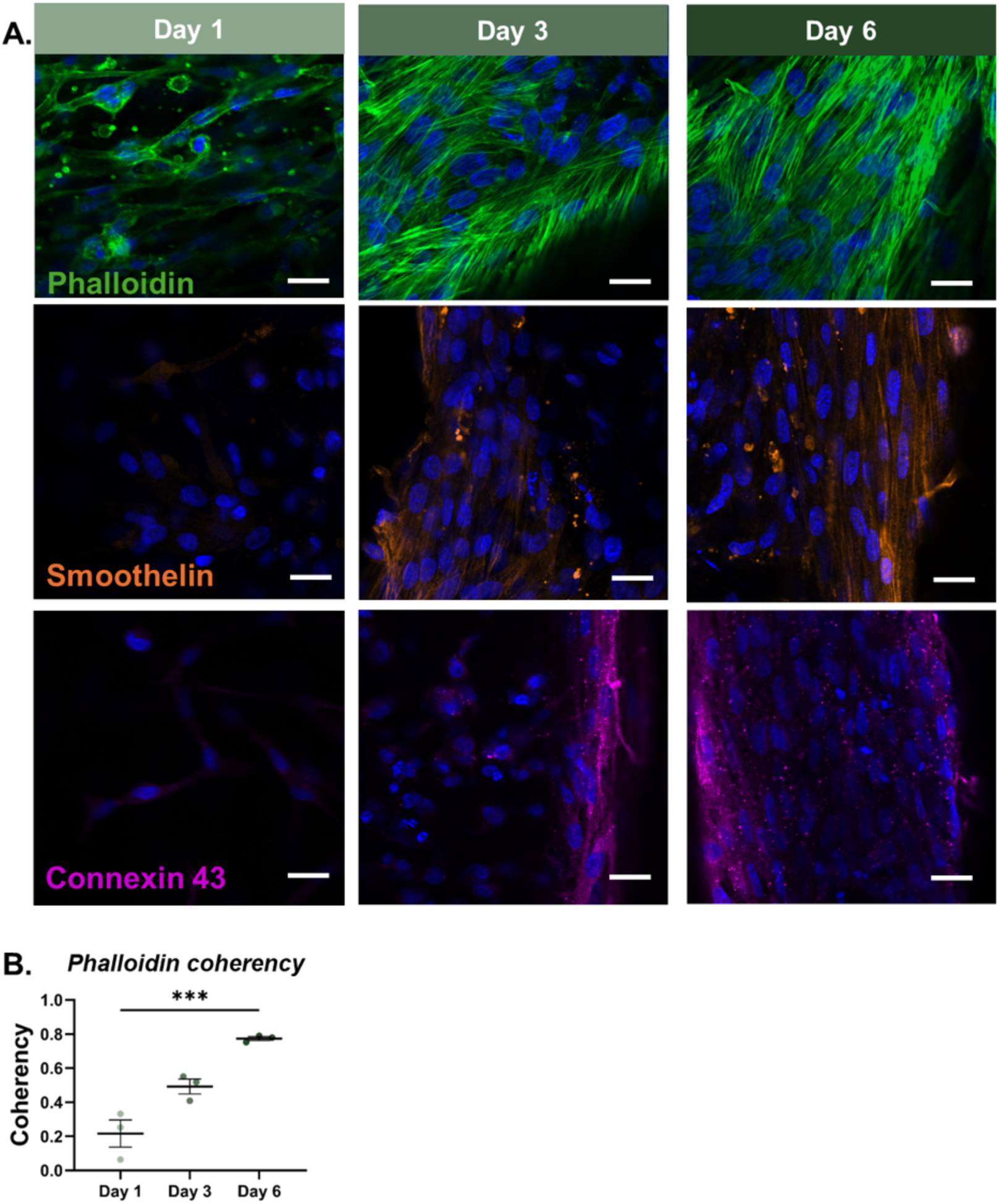
Structural Maturation of OXTR KO Engineered Myometrial Microtissues. To confirm that genetic deletion of the oxytocin receptor does not impair normal microtissue development, OXTR-KO engineered myometrial microtissues were characterized at days 1, 3, and 6 of culture using whole-mount immunofluorescence imaging following the same protocol as wild-type microtissues. (A) Representative whole-mount immunofluorescence images of OXTR KO engineered myometrial microtissues at days 1, 3, and 6, stained for F-actin (phalloidin, green), smoothelin (orange), and connexin-43 (magenta). Nuclei are counterstained with DAPI (blue). Scale bar = 20 µm. (B) Quantification of cytoskeletal coherency across maturation timepoints, indicating progressive cell alignment. Data are presented as mean ± SEM; n = 3. ***p<0.001; one-way ANOVA.

## REFEaRENCES

1. S. Wray, Insights into the uterus. Exp Physiol 92, 621–631 (2007).

2. H. N. Aguilar, B. F. Mitchell, Physiological pathways and molecular mechanisms regulating uterine contractility. Human Reproduction Update 16, 725–744 (2010).

3. A. Evensen, J. M. Anderson, P. Fontaine, Postpartum Hemorrhage: Prevention and Treatment. Am Fam Physician 95, 442–449 (2017).

4. C. A. Grotegut, M. J. Paglia, L. N. Johnson, B. Thames, A. H. James, Oxytocin exposure during labor among women with postpartum hemorrhage secondary to uterine atony. Am J Obstet Gynecol 204, 56.e51–56 (2011).

5. J. Belghiti, G. Kayem, C. Dupont, R. C. Rudigoz, M. H. Bouvier-Colle, C. Deneux-Tharaux, Oxytocin during labour and risk of severe postpartum haemorrhage: a population-based, cohort-nested case-control study. BMJ Open 1, e000514 (2011).

6. S. Bernitz, A. P. Betran, N. Gunnes, J. Zhang, E. Blix, P. Øian, T. M. Eggebø, R. Dalbye, Association of oxytocin augmentation and duration of labour with postpartum haemorrhage: A cohort study of nulliparous women. Midwifery 123, 103705 (2023).

7. S. Phaneuf, G. Asbóth, M. P. Carrasco, B. R. Liñares, T. Kimura, A. Harris, A. L. Bernal, Desensitization of oxytocin receptors in human myometrium. Hum Reprod Update 4, 625–633 (1998).

8. C. Robinson, R. Schumann, P. Zhang, R. C. Young, Oxytocin-induced desensitization of the oxytocin receptor. Am J Obstet Gynecol 188, 497–502 (2003).

9. M. J. Luckas, S. Wray, A comparison of the contractile properties of human myometrium obtained from the upper and lower uterine segments. Bjog 107, 1309–1311 (2000).

10. S. Arrowsmith, Human Myometrial Contractility Assays. Methods Mol Biol 2384, 29–42 (2022).

11. M. Balki, M. Erik-Soussi, J. Kingdom, J. C. Carvalho, Oxytocin pretreatment attenuates oxytocin-induced contractions in human myometrium in vitro. Anesthesiology 119, 552–561 (2013).

12. M. Balki, M. Erik-Soussi, N. Ramachandran, J. Kingdom, J. C. A. Carvalho, The Contractile Effects of Oxytocin, Ergonovine, and Carboprost and Their Combinations: An In Vitro Study on Human Myometrial Strips. Anesthesia & Analgesia 120, 1074–1084 (2015).

13. R. C. Young, Myocytes, Myometrium, and Uterine Contractions. Annals of the New York Academy of Sciences 1101, 72–84 (2007).

14. C. Busch, C. J. Hill, K. Paterson, R. Mellin, M. Zagnoni, D. K. Hapangama, M. E. Sandison, Functional, patient-derived 3D tri-culture models of the uterine wall in a microfluidic array. Hum Reprod 39, 2537–2550 (2024).

15. A. P. Maxey, M. L. McCain, Tools, techniques, and future opportunities for characterizing the mechanobiology of uterine myometrium. Exp Biol Med (Maywood) 246, 1025–1035 (2021).

16. J. Condon, S. Yin, B. Mayhew, R. A. Word, W. E. Wright, J. W. Shay, W. E. Rainey, Telomerase Immortalization of Human Myometrial Cells1. Biology of Reproduction 67, 506–514 (2002).

17. M. S. Soloff, Y. J. Jeng, M. Ilies, S. L. Soloff, M. G. Izban, T. G. Wood, N. I. Panova, G. V. Velagaleti, G. D. Anderson, Immortalization and characterization of human myometrial cells from term-pregnant patients using a telomerase expression vector. Mol Hum Reprod 10, 685–695 (2004).

18. H. Han, X. Ying, Q. Chen, J. Fang, D. Xu, X. Lyu, J. Zheng, L. Zou, Q. Luo, N. Hu, Monitoring of inflammatory preterm responses via myometrial cell based multimodal electrophysiological and optical biosensing platform. Biosens Bioelectron 274, 117197 (2025).

19. S. Siricilla, K. M. Knapp, J. H. Rogers, C. Berger, E. L. Shelton, D. Mi, P. Vinson, J. Condon, B. C. Paria, J. Reese, Q. Sheng, J. L. Herington, Comparative analysis of myometrial and vascular smooth muscle cells to determine optimal cells for use in drug discovery. Pharmacol Res 146, 104268 (2019).

20. J. Fitzgibbon, J. J. Morrison, T. J. Smith, M. O’Brien, Modulation of human uterine smooth muscle cell collagen contractility by thrombin, Y-27632, TNF alpha and indomethacin. Reprod Biol Endocrinol 7, 2 (2009).

21. A. P. Maxey, J. M. Travis, M. L. McCain, Regulation of oxytocin-induced calcium transients and gene expression in engineered myometrial tissues by tissue architecture and matrix rigidity. Curr Res Physiol 6, 100108 (2023).

22. E. Dallot, M. Pouchelet, N. Gouhier, D. Cabrol, F. Ferré, M. Breuiller-Fouché, Contraction of cultured human uterine smooth muscle cells after stimulation with endothelin-1. Biol Reprod 68, 937–942 (2003).

23. M. J. Taggart, S. Wray, Contribution of sarcoplasmic reticular calcium to smooth muscle contractile activation: gestational dependence in isolated rat uterus. J Physiol 511 (Pt 1), 133–144 (1998).

24. C. Ulrich, K. Siddiqui, L. K. Baldwin, W. Hua, J. K. Kuklok, J. J. Okaikoi, L. L. Parker, J. Petereit, D. R. Quilici, G. M. Silva, A. Sivakoses, J. S. Wong-Fortunato, R. J. Woolsey, A. West, Y. Jin, H. Burkin, A bioprinted model of pregnant human uterine myometrium. Front Bioeng Biotechnol 13, 1632320 (2025).

25. G. R. Souza, H. Tseng, J. A. Gage, A. Mani, P. Desai, F. Leonard, A. Liao, M. Longo, J. S. Refuerzo, B. Godin, Magnetically Bioprinted Human Myometrial 3D Cell Rings as A Model for Uterine Contractility. Int J Mol Sci 18, (2017).

26. P. Niessen, S. Rensen, J. van Deursen, J. De Man, A. De Laet, J. M. Vanderwinden, T. Wedel, D. Baker, P. Doevendans, M. Hofker, M. Gijbels, G. van Eys, Smoothelin-a is essential for functional intestinal smooth muscle contractility in mice. Gastroenterology 129, 1592–1601 (2005).

27. F. T. van der Loop, G. Schaart, E. D. Timmer, F. C. Ramaekers, G. J. van Eys, Smoothelin, a novel cytoskeletal protein specific for smooth muscle cells. J Cell Biol 134, 401–411 (1996).

28. K. Saurabh, M. N. Mbadhi, K. K. Prifti, K. T. Martin, A. I. Frolova, Sphingosine 1-Phosphate Activates S1PR3 to Induce a Proinflammatory Phenotype in Human Myometrial Cells. Endocrinology 164, bqad066 (2023).

29. S. Arrowsmith, P. Keov, M. Muttenthaler, C. W. Gruber, Contractility Measurements of Human Uterine Smooth Muscle to Aid Drug Development. J Vis Exp, (2018).

30. J. R. G. Challis, S. G. Matthews, W. Gibb, S. J. Lye, Endocrine and Paracrine Regulation of Birth at Term and Preterm*. Endocrine Reviews 21, 514–550 (2000).

31. M. Malik, M. Roh, S. K. England, Uterine contractions in rodent models and humans. Acta Physiol (Oxf) 231, e13607 (2021).

32. S. Weiss, T. Jaermann, P. Schmid, P. Staempfli, P. Boesiger, P. Niederer, R. Caduff, M. Bajka, Three-dimensional fiber architecture of the nonpregnant human uterus determined ex vivo using magnetic resonance diffusion tensor imaging. Anat Rec A Discov Mol Cell Evol Biol 288, 84–90 (2006).

33. Y.-W. Chan, H. A. van den Berg, J. D. Moore, S. Quenby, A. M. Blanks, Assessment of myometrial transcriptome changes associated with spontaneous human labour by high-throughput RNA-seq. Experimental Physiology 99, 510–524 (2014).

34. O. Shynlova, P. Tsui, S. Jaffer, S. J. Lye, Integration of endocrine and mechanical signals in the regulation of myometrial functions during pregnancy and labour. European Journal of Obstetrics & Gynecology and Reproductive Biology 144, S2–S10 (2009).

35. G. K. Owens, Regulation of differentiation of vascular smooth muscle cells. Physiol Rev 75, 487–517 (1995).

36. G. K. Owens, M. S. Kumar, B. R. Wamhoff, Molecular regulation of vascular smooth muscle cell differentiation in development and disease. Physiol Rev 84, 767–801 (2004).

37. M. T. Lam, Y. C. Huang, R. K. Birla, S. Takayama, Microfeature guided skeletal muscle tissue engineering for highly organized 3-dimensional free-standing constructs. Biomaterials 30, 1150–1155 (2009).

38. K. R. Stevens, K. L. Kreutziger, S. K. Dupras, F. S. Korte, M. Regnier, V. Muskheli, M. B. Nourse, K. Bendixen, H. Reinecke, C. E. Murry, Physiological function and transplantation of scaffold-free and vascularized human cardiac muscle tissue. Proc Natl Acad Sci U S A 106, 16568–16573 (2009).

39. C. Lui, A. F. Chin, S. Park, E. Yeung, C. Kwon, G. Tomaselli, Y. Chen, N. Hibino, Mechanical stimulation enhances development of scaffold-free, 3D-printed, engineered heart tissue grafts. J Tissue Eng Regen Med 15, 503–512 (2021).

40. T. Kimura, O. Tanizawa, K. Mori, M. J. Brownstein, H. Okayama, Structure and expression of a human oxytocin receptor. Nature 356, 526–529 (1992).

41. B. M. Sanborn, C. Y. Ku, S. Shlykov, L. Babich, Molecular signaling through G-protein-coupled receptors and the control of intracellular calcium in myometrium. J Soc Gynecol Investig 12, 479–487 (2005).

42. G. Chioss, M. M. Costantine, E. Bytautiene, A. Betancourt, G. D. Hankins, G. R. Saade, M. Longo, In vitro myometrial contractility profiles of different pharmacological agents used for induction of labor. Am J Perinatol 29, 699–704 (2012).

43. J. E. Gullam, A. M. Blanks, S. Thornton, A. Shmygol, Phase-plot analysis of the oxytocin effect on human myometrial contractility. European Journal of Obstetrics & Gynecology and Reproductive Biology 144, S20–S24 (2009).

44. A. Budden, L. J. Chen, A. Henry, High-dose versus low-dose oxytocin infusion regimens for induction of labour at term. Cochrane Database Syst Rev 2014, Cd009701 (2014).

45. ACOG Practice Bulletin No. 107: Induction of labor. Obstet Gynecol 114, 386–397 (2009).

46. D. Daly, K. C. S. Minnie, A. Blignaut, E. Blix, A. B. Vika Nilsen, A. Dencker, K. Beeckman, M. M. Gross, J. Pehlke-Milde, S. Grylka-Baeschlin, M. Koenig-Bachmann, J. A. Clausen, E. Hadjigeorgiou, S. Morano, L. Iannuzzi, B. Baranowska, I. Kiersnowska, K. Uvnäs-Moberg, How much synthetic oxytocin is infused during labour? A review and analysis of regimens used in 12 countries. PLoS One 15, e0227941 (2020).

47. S. Wray, Uterine contraction and physiological mechanisms of modulation. Am J Physiol 264, C1–18 (1993).

48. S. Wray, S. Arrowsmith, Uterine Excitability and Ion Channels and Their Changes with Gestation and Hormonal Environment. Annu Rev Physiol 83, 331–357 (2021).

49. S. Wray, T. Burdyga, D. Noble, K. Noble, L. Borysova, S. Arrowsmith, Progress in understanding electro-mechanical signalling in the myometrium. Acta Physiol (Oxf) 213, 417–431 (2015).

50. M. Balki, N. Ramachandran, S. Lee, C. Talati, The Recovery Time of Myometrial Responsiveness After Oxytocin-Induced Desensitization in Human Myometrium In Vitro. Anesth Analg 122, 1508–1515 (2016).

51. F. Conti, S. Sertic, A. Reversi, B. Chini, Intracellular trafficking of the human oxytocin receptor: evidence of receptor recycling via a Rab4/Rab5 “short cycle”. American Journal of Physiology-Endocrinology and Metabolism 296, E532–E542 (2009).

52. B. Jurek, I. D. Neumann, The Oxytocin Receptor: From Intracellular Signaling to Behavior. Physiol Rev 98, 1805–1908 (2018).

53. M. Huang, X. Li, P. Guo, Z. Yu, Y. Xu, Z. Wei, The abnormal expression of oxytocin receptors in the uterine junctional zone in women with endometriosis. Reprod Biol Endocrinol 15, 1 (2017).

54. E. E. Don, V. Mijatovic, J. A. F. Huirne, Infertility in patients with uterine fibroids: a debate about the hypothetical mechanisms. Hum Reprod 38, 2045–2054 (2023).

55. O. N. Richter, K. Tschubel, J. Schmolling, M. Kupka, U. Ulrich, E. Wardelmann, Immunohistochemical reactivity of myometrial oxytocin receptor in extracorporeally perfused nonpregnant human uteri. Arch Gynecol Obstet 269, 16–24 (2003).

56. D. Y. Kim, J. M. Reynaud, A. Rasalouskaya, I. Akhrymuk, J. A. Mobley, I. Frolov, E. I. Frolova, New World and Old World Alphaviruses Have Evolved to Exploit Different Components of Stress Granules, FXR and G3BP Proteins, for Assembly of Viral Replication Complexes. PLOS Pathogens 12, e1005810 (2016).

57. C. A. Collier, K. I. Ortega Sandoval, A. Salikhova, S. Rengarajan, A. Tharakesh, A. A. Millan, S. Srinivasan, S. A. Raghavan, Macrophage Immune-Competent Colon Assembloids for Functional Interrogation of Neuroinflammation-Induced Colonic Dysmotility. Gastro Hep Advances, 100894 (2026).

58. J. Schindelin, I. Arganda-Carreras, E. Frise, V. Kaynig, M. Longair, T. Pietzsch, S. Preibisch, C. Rueden, S. Saalfeld, B. Schmid, J.-Y. Tinevez, D. J. White, V. Hartenstein, K. Eliceiri, P. Tomancak, A. Cardona, Fiji: an open-source platform for biological-image analysis. Nature Methods 9, 676–682 (2012).

59. N. García-Quintáns, S. Sacristán, C. Márquez-López, C. Sánchez-Ramos, F. Martinez-de-Benito, D. Siniscalco, A. González-Guerra, E. Camafeita, M. Roche-Molina, M. Lytvyn, D. Morera, M. I. Guillen, M. A. Sanguino, D. Sanz-Rosa, D. Martín-Pérez, R. Garcia, J. A. Bernal, MYH10 activation rescues contractile defects in arrhythmogenic cardiomyopathy (ACM). Nature Communications 14, 6461 (2023).

60. P. Holder. (2026).

61. M. I. Love, W. Huber, S. Anders, Moderated estimation of fold change and dispersion for RNA-seq data with DESeq2. Genome Biology 15, 550 (2014).

62. S. Xu, E. Hu, Y. Cai, Z. Xie, X. Luo, L. Zhan, W. Tang, Q. Wang, B. Liu, R. Wang, W. Xie, T. Wu, L. Xie, G. Yu, Using clusterProfiler to characterize multiomics data. Nature Protocols 19, 3292–3320 (2024).

